# Potassium deficiency reinforces the endodermal but not the exodermal suberized barrier in maize seminal roots

**DOI:** 10.64898/2026.09.21.753126

**Authors:** Tingting Liu, Klaus Dittert, Ismail Cakmak, Jóska Gerendás, Paul Grünhofer, Viktoria Zeisler-Diehl, Lukas Schreiber, Tino Kreszies

**Affiliations:** Department of Crop Sciences, Division of Plant Nutrition and Crop Physiology, University of Göttingen, Carl-Sprengel-Weg 1, 37075 Göttingen, Germany; Faculty of Engineering and Natural Sciences, Sabancı University, 34956 Istanbul, Turkey; K+S Minerals and Agriculture GmbH, Bertha-von-Suttner-Str. 7, 34131 Kassel, Germany; Department of Ecophysiology, Institute of Cellular and Molecular Botany, University of Bonn, Kirschallee 1, 53115 Bonn, Germany; Centre for Crop Systems Analysis, Wageningen University and Research, Wageningen, the Netherlands

**Keywords:** abscisic acid (ABA), apoplastic barrier, endodermis, exodermis, K deficiency, maize, rubidium flux, suberin

## Abstract

Potassium (K) deficiency is a widespread constraint on maize production, yet how it influences the suberized apoplastic barriers that control radial K transport in roots, and whether the endodermis and the constitutively suberized exodermis respond differently, has remained unclear. We grew maize under K deficiency in soil and across a hydroponic K gradient, with abscisic acid (ABA) and fluridone treatments, and analysed seminal roots by Fluorol Yellow 088 staining, tissue-resolved suberin chemistry of endodermis and exodermis, rubidium (Rb⁺) flux, and RNA-sequencing with weighted gene co-expression network analysis (WGCNA). K deficiency selectively increased endodermal aliphatic suberin and eliminated endodermal passage cells in soil-grown roots, whereas exodermal suberin was unchanged. Although K⁺ influx remained high, root-to-shoot Rb⁺ translocation fell sharply, consistent with K retention. A single co-expression module linked the suberin-biosynthetic genes with several K transport genes, and the suberin programme responded to exogenous ABA, indicating ABA-dependent co-regulation. Soil and hydroponic systems converged on the same endodermis-specific anatomical response. Endodermal, but not exodermal, barrier reinforcement limits K leakage from the stele in maize roots. This tissue-specific plasticity identifies the endodermal suberin biosynthesis programme, and its coordination with K⁺ uptake, as a target for improving K use efficiency.

**Highlight:** Potassium deficiency reinforces the endodermal but not the exodermal suberin barrier in maize roots, restricting root-to-shoot K translocation through an ABA-dependent programme co-regulated with K transport.

## Introduction

Potassium (K) is the most abundant cation in plant cells and an essential macronutrient that plays a central role in enzyme activation, photosynthesis, osmoregulation, turgor-driven growth, stomatal movement, the long-distance transport of photoassimilates, and cellular charge balance (Zörb et al., 2014; Hawkesford et al., 2023). Potassium also plays a critical role in crop tolerance to abiotic and biotic stress, partly by limiting the accumulation of reactive oxygen species. Consequently, K-deficient plants become more susceptible to drought, salinity, low temperature, ion toxicity and pathogen attack (Cakmak, 2005). Although total soil K reserves are usually large, the readily plant-available fraction is often small. Crops therefore depend on the slow release of K from soil minerals and on K fertilization, for which global demand continues to rise. Crop species differ markedly in their K demand (Sardans & Peñuelas, 2021). Maize has a high K requirement, taking up large amounts of K during vegetative growth and relying on a diverse set of K channels and transporters for uptake and remobilization (Coskun et al., 2014; Nieves-Cordones et al., 2014; Lyzenga et al., 2023). An inadequate or inefficient K supply therefore readily constrains growth and yield (Zhang et al., 2023). Because much of the crop is grown on soils where readily available K is limited, the efficiency with which maize roots acquire and retain K is of both physiological and agronomic interest.

K taken up at the root surface must cross the root radially to reach the xylem. According to the composite transport model, this radial movement of water and solutes follows three parallel pathways: the apoplastic route through the cell walls, the symplastic route through plasmodesmata, and the transcellular route across successive membranes (Steudle & Peterson, 1998; Ranathunge et al., 2017; Robe & Barberon, 2023). The apoplastic route is controlled by two cell layers that deposit specialized wall modifications, the endodermis and the exodermis, and in maize the endodermis is the principal barrier to the radial movement of ions (Peterson et al., 1993). The endodermis first forms a Casparian strip, a band of lignin in the anticlinal walls that seals the apoplast and forces solutes across the selective endodermal plasma membrane (Schreiber et al., 1999; Geldner, 2013). Subsequently, the endodermis deposits suberin lamellae over the whole cell surface, a secondary wall modification that further restricts transmembrane uptake and the back-diffusion of solutes (Schreiber et al., 1999; Kreszies et al., 2018). Individual endodermal cells can nevertheless remain unsuberized as passage cells, retaining only the Casparian strip and providing localized low-resistance sites for exchange between the cortex and the stele (Kreszies et al., 2018).

Suberin is a glycerol-based polyester with aliphatic and aromatic domains, deposited as lamellae between the plasma membrane and the primary cell wall. Its synthesis draws on fatty acid elongation, ω-hydroxylation, and aromatic acylation (Franke & Schreiber, 2007; Pollard et al., 2008). The quantity of suberin laid down in the endodermis depends on genotype and environment interactions: under stress it can increase or, where suberin biosynthesis is down-regulated, be lower compared to controls. In crop roots, abiotic stresses including drought, osmotic, and waterlogging stress markedly increase endodermal suberization, as documented in barley seminal roots and rice adventitious roots (Ranathunge et al., 2011; Kreszies et al., 2019). This response is associated with the phytohormone abscisic acid (ABA) and, based on work in Arabidopsis, is mediated by a set of MYB transcription factors that control suberin biosynthetic genes (Shukla et al., 2021; Meng et al., 2025). Nutrient supply likewise modulates suberization, increasing endodermal suberin under K and sulphur deficiency in Arabidopsis (Barberon et al., 2016; Barberon, 2017) and altering it according to nutrient status in crop roots (Grünhofer et al., 2021). Such cues are not universal, however: iron deficiency reduces root suberin in dicots but not in maize (Sijmons et al., 1985). Even in Arabidopsis the link between suberin and the transport of any single nutrient is not straightforward. Across the many suberin- and Casparian-strip-altered mutants surveyed by Grünhofer et al. (2024), leaf concentrations of individual ions, including K, respond inconsistently, and the effects of the suberin lamellae are hard to separate from those of the Casparian strip. The role of endodermal suberin in nutrient transport therefore remains to be established directly in crops.

Cereal roots differ from the Arabidopsis model in possessing a second apoplastic barrier, the exodermis. This suberized cell layer beneath the rhizodermis forms an additional, outer checkpoint to radial transport and is found in most angiosperms, Arabidopsis being one of the exceptions that lack it (Hose et al., 2001; Liu & Kreszies, 2023). In maize the exodermis is suberized already under non-stressed conditions (Schreiber et al., 1999; Kreszies et al., 2018; Liu et al., 2026), so that the root presents two suberized barriers that could in principle be regulated together or independently. These barriers can respond differently to ABA, and the response appears to depend on root type. In the adventitious roots of rice and barley, ABA promotes exodermal or hypodermal suberization and the formation of a barrier to radial oxygen loss under waterlogging (Shiono et al., 2022; Shiono & Matsuura, 2024). In barley seminal roots, by contrast, exogenous ABA promotes endodermal but not hypodermal suberization (Grünhofer et al., 2021). With its dimorphic complement of a constitutive exodermis and a developmentally regulated endodermis, the maize seminal root is well suited to ask which barrier responds to a nutrient stress, and how.

For nutrient deficiencies, the remodelling of these barriers in maize has so far been characterized only for nitrogen deficiency, which reinforced the endodermis more than the exodermis (Liu et al., 2026). How K deficiency affects the barriers, and what this means for K transport, has not yet been examined. A transcriptomic survey of maize roots under K deficiency revealed extensive changes in K signalling and transporter genes (Guo et al., 2023), but the accompanying anatomical and biochemical changes in the apoplastic barriers, and their link to K movement, were not addressed. Suberin is increasingly implicated in controlling the radial movement of solutes into and out of the stele. In Arabidopsis and barley, more heavily suberized root zones translocate K more efficiently to the shoot and lose less of a caesium tracer used as a K analogue (Vestenaa et al., 2024). In suberin-deficient poplar, reduced endodermal suberin increased the inward bypass of sodium chloride and the uptake and translocation of a herbicide tracer (Grünhofer et al., 2024). Because the maize exodermis is already suberized and putatively forms an outer control point for K entry, it was not evident a priori whether K deficiency would further reinforce both barriers or act selectively on either of them. We therefore investigated whether and how K deficiency remodels the endodermal and exodermal barriers of maize seminal roots, and whether any reinforcement is linked to K retention within the root. We combined soil- and hydroponically grown maize with anatomical staining, biochemical quantification of endodermal and exodermal suberin, transcriptome sequencing with weighted gene co-expression network analysis, and rubidium influx and translocation measurements as a proxy for K transport. To test whether ABA contributes to the response, we applied exogenous ABA and the ABA-biosynthesis inhibitor fluridone.

## Materials and Methods

### Plant material and growth conditions

Maize (cv. Ronaldinio; KWS Saat SE & Co. KGaA, Einbeck, Germany) was used throughout all experiments. Ronaldinio is a commercially available, early-maturing silage maize cultivar widely grown under central European conditions and has previously been used to study root suberization under nitrogen deficiency (Liu et al., 2026). In both experiments, seedlings were raised in controlled-environment chambers (23/18 °C day/night, 14/10 h photoperiod, 400 μmol photons m⁻Z s⁻¹, 65/75% relative humidity day/night). Uniform seedlings of comparable size were selected on day 6 after germination for transplanting. Germination differed between the two experiments and is described below.

### Soil experiment

#### Experimental design and plant cultivation

Soil was collected from an agricultural site in Rudolstadt-Schwarza, Germany (50°41′27.6″N, 11°19′46.1″E) and mixed with quartz sand at a 5:3 (w/w) ratio to yield a K concentration of 29 mg K kg⁻¹ dry weight for the K-deficient treatment (−K). For the K-sufficient control (+K), K₂SO₄ was incorporated to raise the K concentration to 100 mg K kg⁻¹ dry weight. These levels correspond to soil K classes A (deficient) and C (adequate) of the German fertilizer recommendation scheme (VDLUFA, 1998). Soil texture, pH, and nutrient status were characterized prior to the experiment (Table S1). Water-holding capacity was determined gravimetrically by saturating soil columns for 24 h, draining for 48 h, and recording water retention. Soil moisture was logged throughout cultivation (Fig. S1). Soil volumetric water content was maintained at 80% of maximum water-holding capacity by daily watering with deionized water and monitored using capacitance sensors (EC-5, Decagon Devices, Pullman, WA, USA) installed at 25% and 75% of column depth.

For the soil experiment, seeds were germinated directly in soil-filled cups (10 cm depth) for 5 d. Cylindrical PVC columns (120 cm × 15 cm inner diameter) lined with polyethylene film were filled with 28 kg of the respective soil mixture, and six-day-old seedlings were transplanted into the columns (13 plants for +K and 12 for −K) and grown for 25 d, giving a total plant age of 31 d at harvest. At harvest, shoots were excised and the root system was carefully recovered. Seminal roots were identified and a segment at approximately 6% of total root length from the root tip, a position defined on the basis of Fluorol Yellow 088 staining (see below), was immediately snap-frozen in liquid nitrogen for RNA extraction. The remaining root material was used for anatomical, biochemical, and elemental analyses.

### Hydroponic experiment

#### Experimental design and plant cultivation

For the hydroponic experiment, seeds were surface-sterilized in 2% (v/v) NaOCl for 4 min, rinsed thoroughly with deionized water, and germinated on moist paper rolls soaked in 1 mM CaSO₄ in darkness at 23 °C for 2 d, followed by a further 3 d in the controlled-environment chambers. Six-day-old seedlings were then transferred to quarter-strength nutrient solution for 4 d before transfer to full-strength nutrient solution. The full-strength base solution contained (in μM): 2000 Ca(NO₃)₂, 750 MgSO₄, 100 Ca(H₂PO₄)₂, 50 CaCl₂, 100 Fe-EDTA, 1 H₃BO₃, 1.3 MnSO₄, 0.2 CuSO₄, 0.2 ZnSO₄, and 0.01 (NH₄)₆Mo₇O₂₄, without added K. The nutrient solution was renewed every 3 d and adjusted to pH 6.0–6.2 at each change.

Six treatments were established by supplementing the base solution with K₂SO₄ and/or phytohormones: 25 μM K (K25), 100 μM K (K100), 600 μM K (K600, control), 5000 μM K (K5000), 600 μM K + 10 μM abscisic acid (K600+ABA), and 600 μM K + 10 μM fluridone (K600+FLU). The concentration of 600 μM K was selected as the control based on preliminary growth experiments. Exogenous ABA at 10 μM was applied to stimulate suberin deposition in endodermis and exodermis (Shiono et al., 2022; Grünhofer et al., 2021). Fluridone, an inhibitor of ABA biosynthesis (Gamble & Mullet, 1986), was applied at 10 μM to suppress suberization, following its use at comparable concentrations in rice adventitious roots (Shiono et al., 2022). Phytohormones were replenished with each solution change.

Plants were grown in 5 L opaque containers with three plants per container. Five containers were used per treatment, and the three plants of a container were pooled and treated as one biological replicate (n = 5). Shoots and roots were harvested separately 15 d after the onset of full-strength treatment, corresponding to a total plant age of 25 d. Shoot and root dry weights were determined after oven-drying at 60 °C for 72 h.

### Root anatomy and suberin staining

Seminal root segments were collected at defined positions along the root axis, expressed as a percentage of total seminal root length from the apex (defined as 0%). For soil-grown plants, cross-sections were taken at 1%, 12% and 25% for imaging of suberin lamella development. For hydroponically grown plants, cross-sections were taken at 25%, 50% and 70% for the same purpose.

Root segments were either used fresh or fixed in 3.7% (v/v) formaldehyde in phosphate-buffered saline. Transverse sections of 30 μm thickness were prepared using a cryomicrotome (Kedi, China) at −25 °C. Suberin lamellae were visualized by staining sections with 0.01% (w/v) Fluorol Yellow 088 for 1 h in the dark (Brundrett et al., 1991), followed by rinsing with deionized water. Sections were imaged under UV excitation using a fluorescence microscope (Carl Zeiss Microscopy, Jena, Germany). Passage cells were identified as endodermal cells lacking Fluorol Yellow 088 fluorescence and counted manually; results are expressed as the proportion of total endodermal cells counted per cross-section.

### Elemental analysis

Oven-dried plant material was finely ground and digested in concentrated HNO₃/H₂O₂ (2:1, v/v) in a closed-vessel microwave digestion system (Ethos.lab, MLS GmbH, Leutkirch, Germany). Digests were diluted to 25 mL with ultrapure water and K concentrations determined by inductively coupled plasma optical emission spectrometry (ICP-OES; Spectro Genesis, SPECTRO Analytical Instruments GmbH, Kleve, Germany). Soil K availability was determined by extracting air-dried soil with calcium lactate/acetate solution (Schüller, 1969) and measuring K by flame photometry (Eppendorf Elex 6361, Hamburg, Germany).

### Suberin biochemical analysis

Biochemical suberin analysis was performed on seminal roots from both soil-grown and hydroponically grown plants. Root zones were defined as a percentage of total root length from the apex and analysed separately, with three biological replicates per treatment: Zone A (0–25%), Zone B (25–50%) and Zone C (50–100%) for soil-grown plants, and Zones A and B for hydroponically grown plants. Cell walls were isolated by enzymatic digestion in a combined solution of 0.5% (w/v) cellulase and 0.5% (w/v) pectinase in 50 mM citrate buffer (pH 3.5) for three weeks at room temperature under continuous agitation, with the enzyme solution replaced four times. Once a distinct outer cell wall layer was visible under stereomicroscopy, cell walls were separated and washed in 10 mM sodium borate buffer for 24 h to remove enzyme residues. Extractable lipids were removed by incubation in chloroform:methanol (1:1, v/v) for two weeks under continuous agitation. Residues were dried on polytetrafluoroethylene plates and stored at room temperature.

Suberin monomers were released by transesterification in BF₃–methanol at 70 °C overnight, extracted into chloroform with 10 µg dotriacontane as internal standard, and derivatized with 20 µl each of BSTFA and pyridine for 45 min at 70 °C. Monomers were quantified by gas chromatography with flame ionization detection (GC-FID) as described by Baales et al. (2021). Identification of individual suberin monomers was achieved on a GC-MS system by comparing their fragmentation patterns with an in-house database library and with patterns described in the literature. Monomer amounts were normalized to the cylindrical surface area of the respective tissue layer (A = 2πrL, where r is the tissue radius, determined microscopically, and L is the zone length).

### Rubidium flux measurements

Root-to-shoot K transport capacity was assessed using Rb⁺ as a K tracer (Beier et al., 2022). On day 14 of hydroponic treatment, and under the same controlled-environment conditions, roots were sequentially transferred to 1 mM CaSO₄ for 1 min to desorb apoplastically bound ions, to 1 mM RbCl for 20 min, and again to 1 mM CaSO₄ for 1 min. Shoots and roots were separated, oven-dried at 60 °C for 72 h, and digested as described for elemental analysis. Rb concentrations were determined by inductively coupled plasma optical emission spectrometry (ICP-OES; Vista-Pro Axial, Varian Pty Ltd, Mulgrave, Australia). Rb⁺ influx and root-to-shoot translocation rates were calculated according to Beier et al. (2022):

Rb influx (μg g⁻¹ root DW min⁻¹) = (A × C + B × D) / C / 20
Root-to-shoot translocation (μg g⁻¹ root DW min⁻¹) = (B × D) / C / 20

where A = root Rb concentration (μg g⁻¹ DW), B = shoot Rb concentration (μg g⁻¹ DW), C = root dry weight (g), and D = shoot dry weight (g); n = 4 biological replicates.

### RNA extraction and transcriptome sequencing

Fluorol Yellow 088 staining was carried out first, and the resulting patterns of suberin lamella development were used to define the root zones subsequently sampled for RNA extraction. For soil-grown plants, a segment at approximately 6% of total seminal root length from the apex was harvested. For hydroponically grown plants, two zones were collected: zone A (10–20% of root length, onset of suberization) and zone B (40–50%, mature suberized zone). These RNA-seq zones lie within the corresponding zones used for suberin analysis, that is, RNA-seq zone A within Zone A (0–25%) and RNA-seq zone B within Zone B (25–50%); they are therefore sub-regions of the latter rather than identical to them. Tissue was immediately snap-frozen in liquid nitrogen, ground to a fine powder, and total RNA was extracted using the RNeasy Universal Mini Kit (Qiagen, Hilden, Germany). RNA quantity and quality were assessed by spectrophotometry and gel electrophoresis. Four biological replicates per treatment were sent to Novogene (Cambridge, UK) for library preparation and 150 bp paired-end sequencing on an Illumina NovaSeq platform. This yielded 386 million raw reads for the soil experiment (8 libraries) and 2217 million for the hydroponic experiment (48 libraries, six treatments × two root zones × four replicates), giving 2603 million reads from 56 libraries in total.

Raw reads were quality-filtered using fastp (Chen et al., 2018) with default parameters. Clean reads were aligned to the maize B73 reference genome (Zm-B73-REFERENCE-NAM-5.0; Ensembl Plants release 57) using HISAT2 (v2.2.1; Kim et al., 2019), with a mean mapping rate of 83.4% across all 56 libraries. Transcripts were assembled against the reference annotation with StringTie (Pertea et al., 2015), which also yielded novel transcripts not present in the reference annotation; these are reported with ‘novel’ identifiers and were retained for differential expression analysis but not for the co-expression analysis, which was restricted to annotated genes. Gene expression was quantified as FPKM. Differentially expressed genes (DEGs) were identified with DESeq2 (Love et al., 2014) using thresholds of |log₂FC| ≥ 1 and adjusted p ≤ 0.05 (Benjamini–Hochberg correction). Gene Ontology (GO) enrichment and KEGG pathway analyses were performed using clusterProfiler (Yu et al., 2012), with all genes tested for differential expression in the respective comparison as background, term sizes restricted to 10–500 genes, and Benjamini–Hochberg correction applied separately within each ontology (adjusted p ≤ 0.05).

Raw RNA-seq sequencing data have been deposited in the NCBI Sequence Read Archive (SRA) under BioProject accession number PRJNA1517815.

### Weighted gene co-expression network analysis

Co-expression network analysis was performed using the WGCNA R package (Langfelder & Horvath, 2008). Genes with mean FPKM < 1 were excluded, and the approximately 6000 genes with the highest median absolute deviation were retained for analysis (5916 genes in zone A and 5931 in zone B). A signed adjacency matrix was constructed using a soft-thresholding power β of 3 (zone A) or 4 (zone B), selected to achieve scale-free network topology (RZ ≥ 0.85). Modules were identified by hierarchical clustering with dynamic tree cutting (minimum module size 50 genes) and merged at an eigengene correlation threshold of 0.75. Module eigengenes were correlated with physiological traits including shoot and root dry weight, root-to-shoot ratio, Rb⁺ influx, root-to-shoot Rb⁺ translocation, root Rb concentration, and endodermal and exodermal total, aliphatic and aromatic suberin content. Correlations were tested for significance by Student asymptotic p-value; modules with |r| > 0.5 and p < 0.05 were taken forward. For each zone, subsequent analyses focused on the co-expression module most strongly associated with the traits of interest. Hub genes were defined as module members with module membership kME ≥ 0.80 and gene significance for endodermal aliphatic suberin |GS| > 0.5 in their assigned module.

### Statistical analysis

All statistical analyses were performed in R (v4.6.0) using the agricolae package. One-way analysis of variance (ANOVA) followed by Tukey’s HSD post-hoc test was applied at α = 0.05; all tests were two-sided. For hydroponic data, two separate one-way ANOVA models were fitted: one comparing K-gradient treatments (K25, K100, K600, K5000) and one comparing hormone-modified treatments against the K600 control (K600, K600+ABA, K600+FLU). Lowercase letters denote significant differences within the K-gradient comparison; uppercase letters within the hormone comparison. All data are presented as means ± standard deviation (SD).

## Results

### K deficiency reduces plant biomass in soil-grown and hydroponic maize

Maize plants grown in K-deficient soil (−K) showed significant reductions in shoot and root biomass compared to K-sufficient controls (+K; Fig. 1a,b and Fig. 2a,c). The root-to-shoot ratio was also significantly lower in −K plants (Fig. 2e), indicating that root growth was more strongly impaired than shoot growth. In the hydroponic experiment, shoot and root dry weight increased progressively with K supply from K25 to K600, with K5000 not differing significantly from K600 (Fig. 1c and Fig. 2b,d). Fluridone treatment (K600+FLU) bleached the shoots, which appeared almost entirely white, and nearly abolished plant growth, while ABA treatment (K600+ABA) also significantly reduced biomass compared to K600 (Fig. 2b,d). The root-to-shoot ratio was significantly higher under K600+ABA and K600+FLU treatments relative to K25 (Fig. 2f).

**Fig. 1.**
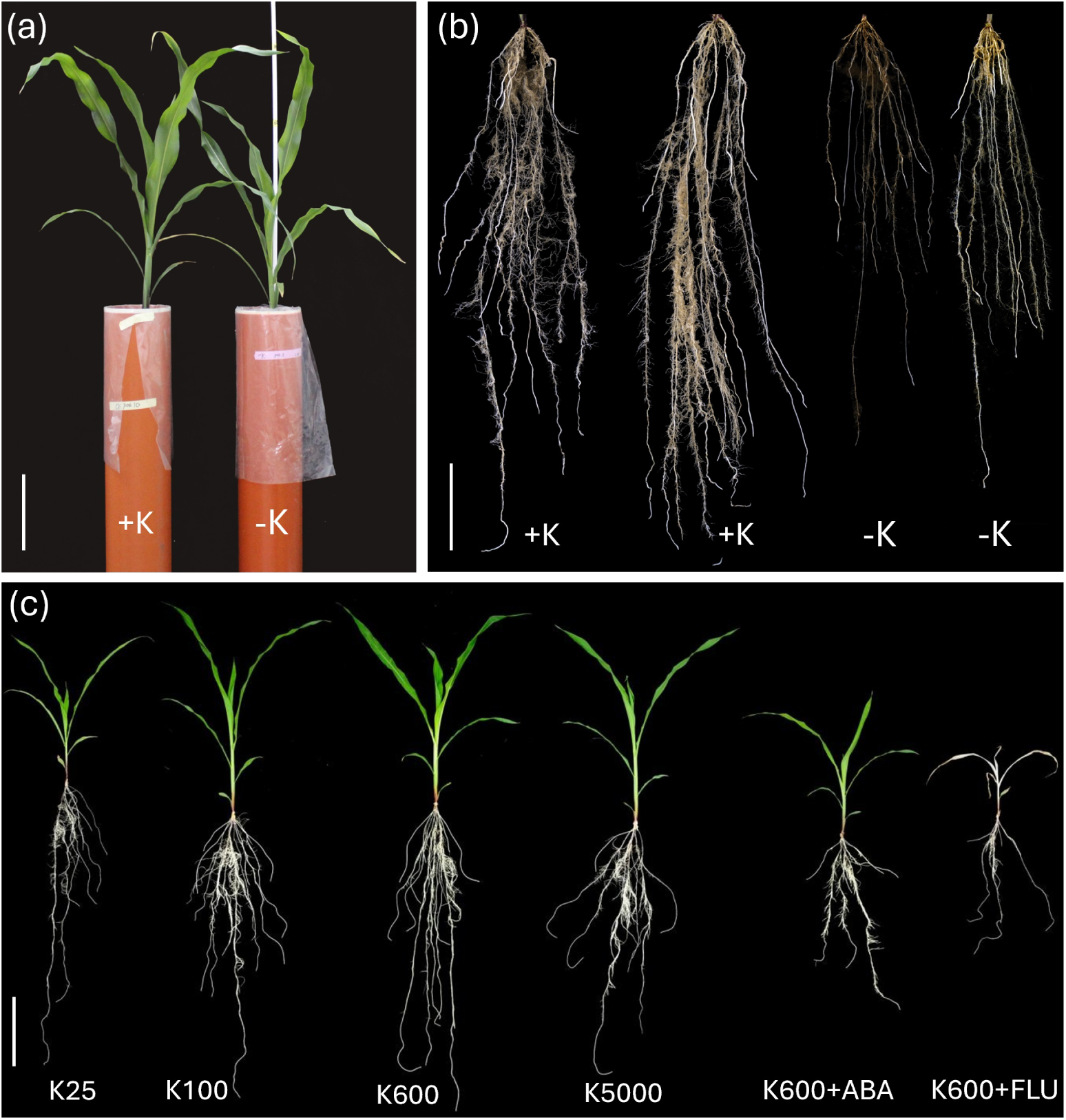
K deficiency reduces growth of soil-grown and hydroponic maize. Representative phenotypes of maize. (a) Shoots of plants grown in K-sufficient (+K) or K-deficient (−K) soil and (b) the excavated seminal root systems (+K and −K). (c) Whole plants grown hydroponically under six K supply and phytohormone treatments (K25, K100, K600, K5000, K600+ABA, K600+FLU). Scale bars, 20 cm. **Alt text:** Photographs of maize plants. Panel a shows two shoots of similar height above pots labelled plus K and minus K. Panel b shows four washed seminal root systems; the two plus K systems are denser and carry more lateral roots than the two minus K systems. Panel c shows six hydroponically grown plants across the K gradient and hormone treatments, with the fluridone plant bleached white and much smaller than the rest.

**Fig. 2.**
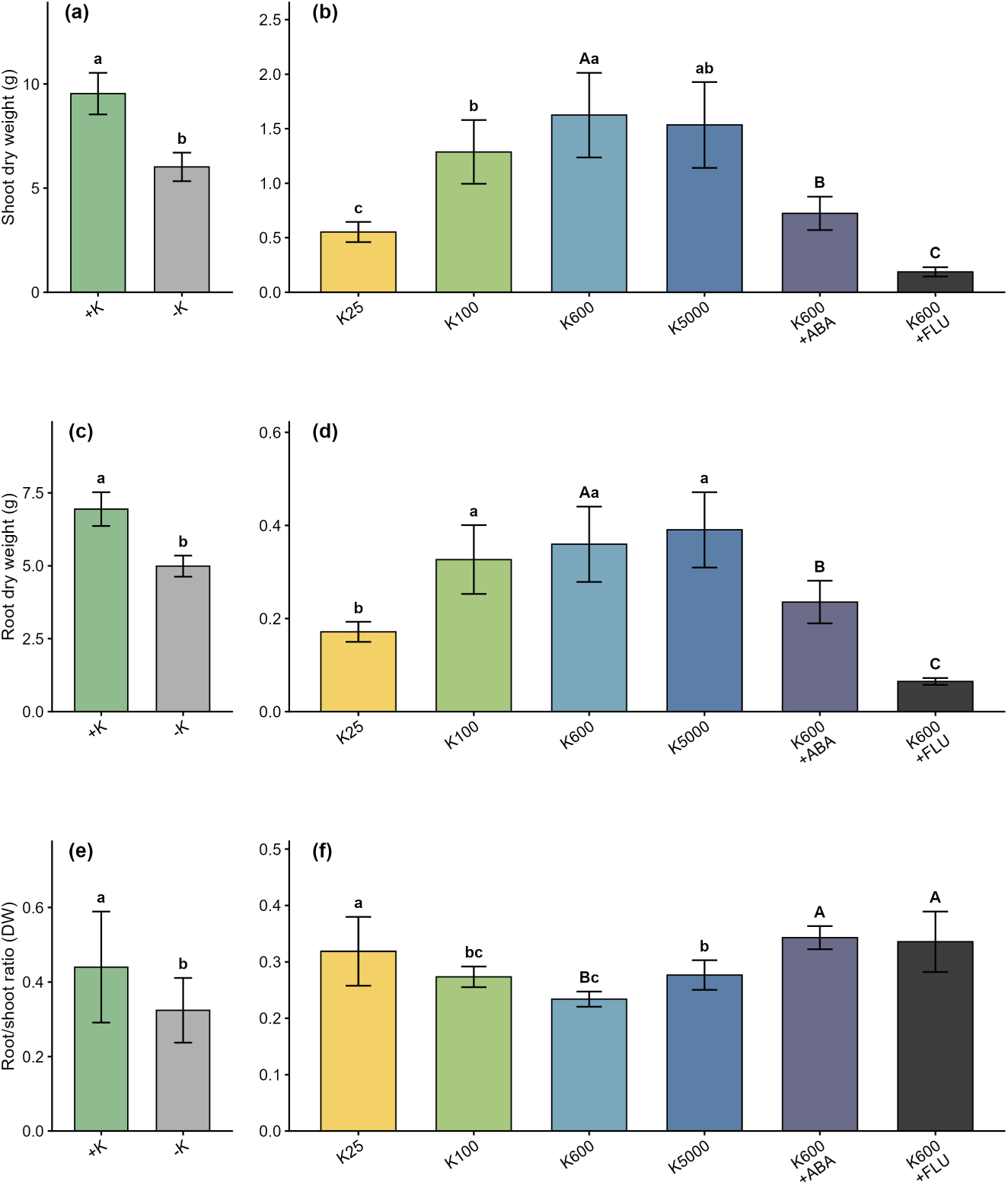
K deficiency reduces shoot and root biomass in soil-grown and hydroponic maize. (a, c, e) Shoot dry weight, root dry weight, and root-to-shoot ratio of plants grown in K-sufficient (+K) or K-deficient (−K) soil. (b, d, f) Shoot dry weight, root dry weight, and root-to-shoot ratio of hydroponically grown plants under six K supply and phytohormone treatments (K25, K100, K600, K5000, K600+ABA, K600+FLU). Bars represent means ± SD (soil, n = 13 for +K and 12 for −K; hydroponic, n = 5 biological replicates). Different letters indicate significant differences at α = 0.05 (Tukey’s HSD, two-sided); for hydroponic data, lowercase letters indicate differences among K-gradient treatments (K25, K100, K600, K5000) and uppercase letters among hormone-modified treatments (K600, K600+ABA, K600+FLU). **Alt text:** Six bar charts of dry weight. The left column compares plus K and minus K soil plants, the right column the six hydroponic treatments; rows show shoot dry weight, root dry weight and root-to-shoot ratio. Bars are means with standard deviation and letters for significant differences. All three measures are lower under minus K, and in hydroponics they increase from K25 to K600 and fall sharply under fluridone.

### K deficiency reduces K concentrations in shoots, roots, and along the root axis

Shoot and root K concentrations were significantly reduced under K deficiency in soil-grown plants (Fig. 3a,b). Analysis of seminal root zones revealed a pronounced basipetal decline in K concentration along the root axis under both treatments, with concentrations highest in Zone A and declining progressively towards Zone C (Fig. 3c). This gradient was significantly attenuated under K deficiency, with lower K concentrations across all zones compared to the +K control (Fig. 3c). In hydroponically grown plants, shoot and root K concentrations increased with increasing K supply across the K-gradient treatments, while K600+FLU plants showed the highest shoot K concentration among all treatments (Fig. 3d,e).

**Fig. 3.**
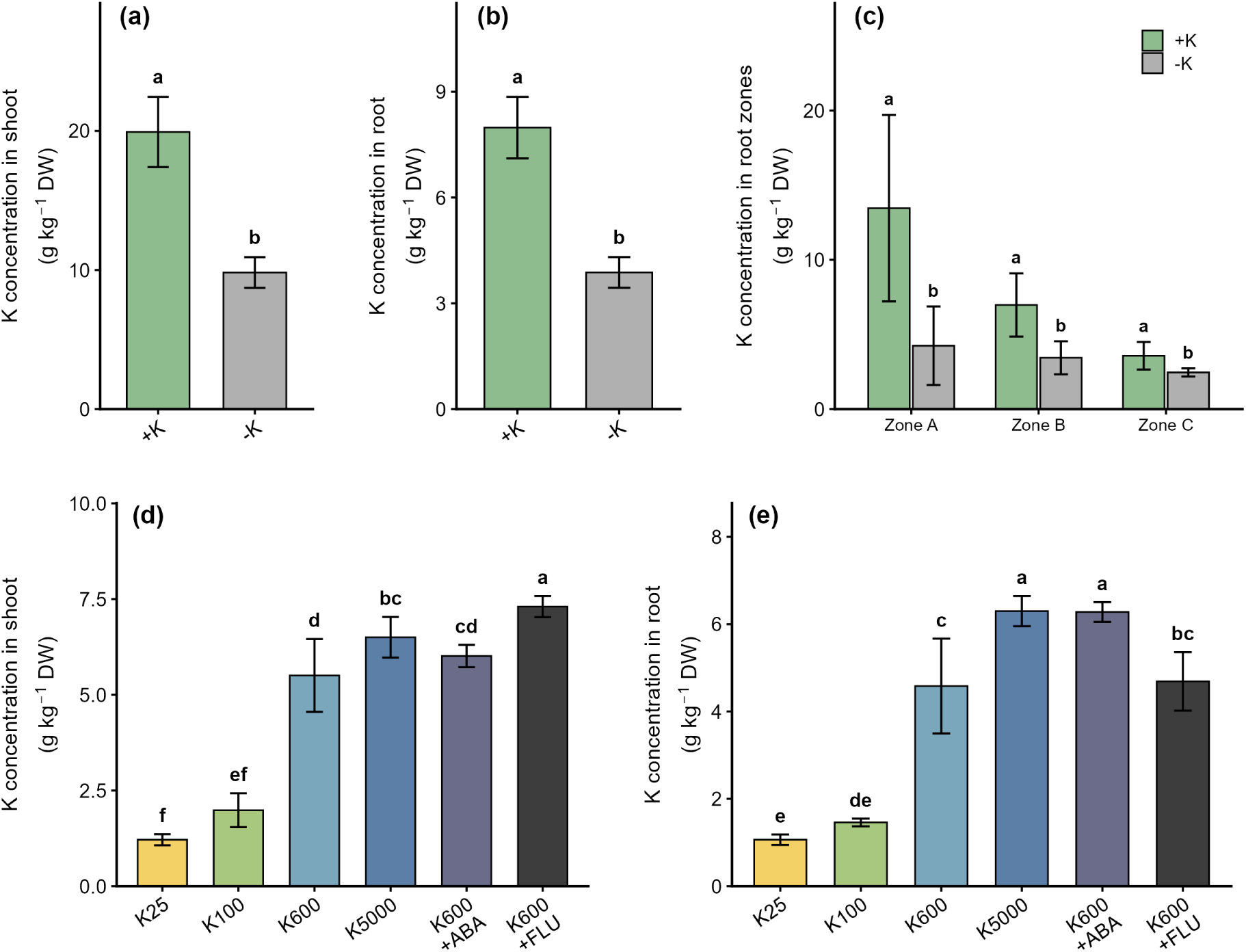
K deficiency lowers potassium concentrations in shoots, roots, and along the seminal root axis. (a, b) K concentration in the shoot and root of plants grown in +K or −K soil. (c) K concentration in three zones of soil-grown seminal roots: Zone A (0–25% of root length from the apex), Zone B (25–50%), and Zone C (50–100%). (d, e) K concentration in the shoot and root of hydroponically grown plants under six K supply and phytohormone treatments. Bars represent means ± SD (soil, n = 11 for +K and 6 for −K in a and b, n = 3 in c; hydroponic, n = 5 biological replicates). Different letters indicate significant differences at α = 0.05 (Tukey’s HSD, two-sided); for hydroponic data, lowercase letters indicate differences among K-gradient treatments and uppercase letters among hormone-modified treatments. **Alt text:** Five bar charts of potassium concentration. Panels a and b give shoot and root concentrations in plus K and minus K soil plants, both markedly lower under minus K. Panel c gives concentrations in root zones A to C, declining from the apex towards the base and lower throughout under minus K. Panels d and e give shoot and root concentrations across the six hydroponic treatments, increasing with K supply.

### K deficiency modulates root-to-shoot K transport capacity

Rb⁺ influx into the root was highest under K100 and slightly higher under K25 compared to K600, indicating upregulation of high-affinity K⁺ uptake systems under K deficiency, and was lowest under K5000 and K600+FLU (Fig. 4a). Root-to-shoot Rb⁺ translocation was highest under K600, substantially exceeding the values under K25 and K100, suggesting greater root retention of K under K deficiency (Fig. 4b). Fluridone dramatically reduced Rb⁺ translocation to near zero, while ABA treatment also significantly reduced translocation relative to K600 (Fig. 4b), consistent with a contribution of ABA to the regulation of root-to-shoot K transport.

**Fig. 4.**
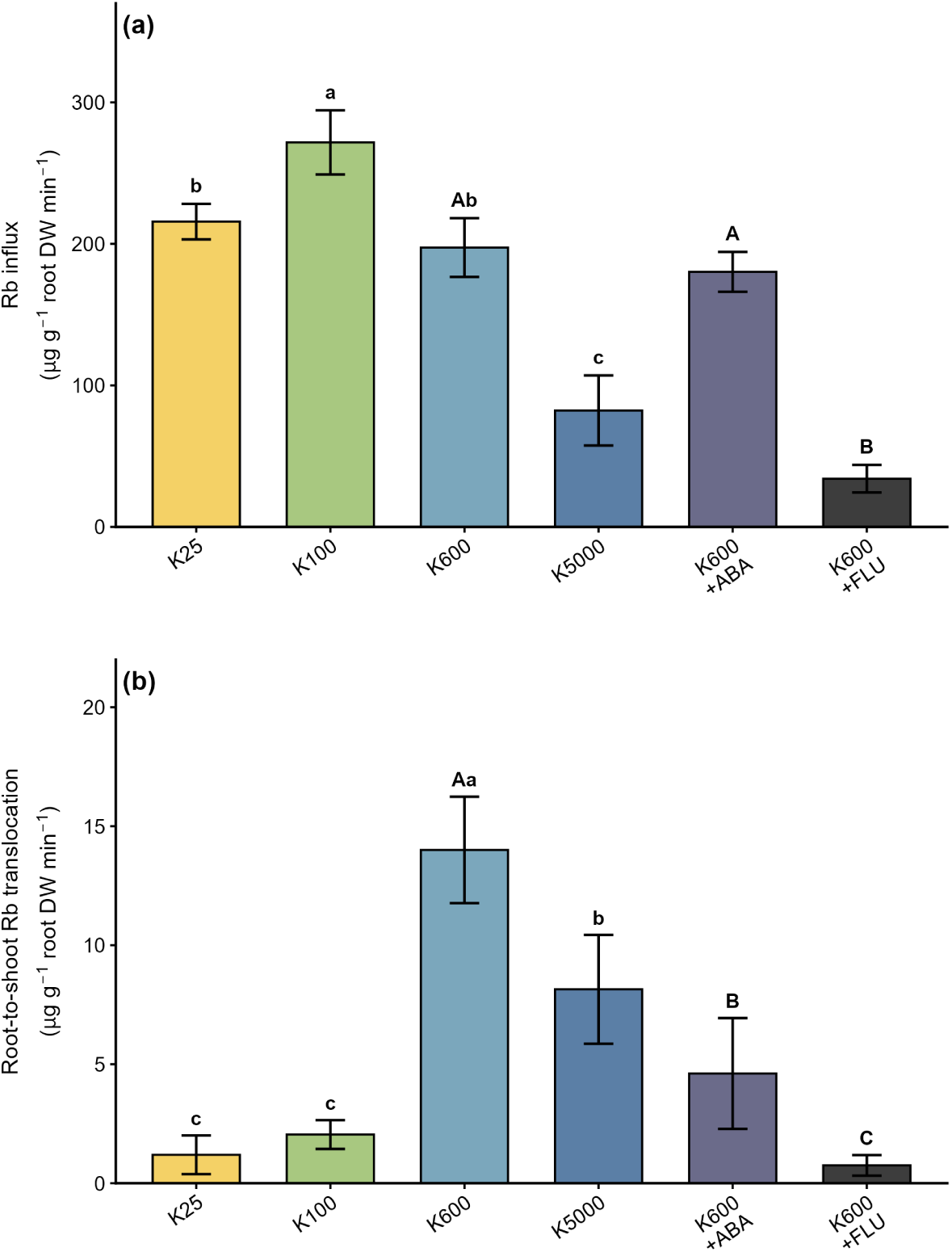
K deficiency modulates root Rb⁺ influx and root-to-shoot Rb⁺ translocation. (a) Rb⁺ influx into roots and (b) root-to-shoot Rb⁺ translocation of hydroponically grown maize under six K supply and phytohormone treatments, measured in a short-term Rb⁺ uptake assay. Rb⁺ was used as a tracer for K⁺ transport. Bars represent means ± SD (n = 4 biological replicates). Different letters indicate significant differences at α = 0.05 (Tukey’s HSD, two-sided); for hydroponic data, lowercase letters indicate differences among K-gradient treatments (K25, K100, K600, K5000) and uppercase letters among hormone-modified treatments (K600, K600+ABA, K600+FLU). **Alt text:** Two bar charts. Panel a shows rubidium influx into roots across six hydroponic treatments, highest under K100 and lowest under K5000 and fluridone. Panel b shows root-to-shoot rubidium translocation, highest under K600 and close to zero under fluridone. Bars are means with standard deviation and letters for significant differences.

### K deficiency selectively increases endodermal aliphatic suberin in soil-grown maize roots

Fluorol Yellow 088 (FY) staining indicated that K deficiency accelerated suberization in seminal roots of soil-grown plants (Fig. 5). Under both conditions, suberin lamellae were absent close to the root tip at 1% relative root length. In the endodermis, continuous suberin lamellae were present at the 12% position in −K plants, whereas suberization was not yet complete in +K plants (Fig. 5b,e). In the exodermis, the FY signal at the 25% position was more pronounced in −K than in +K plants (Fig. 5i,l).

**Fig. 5.**
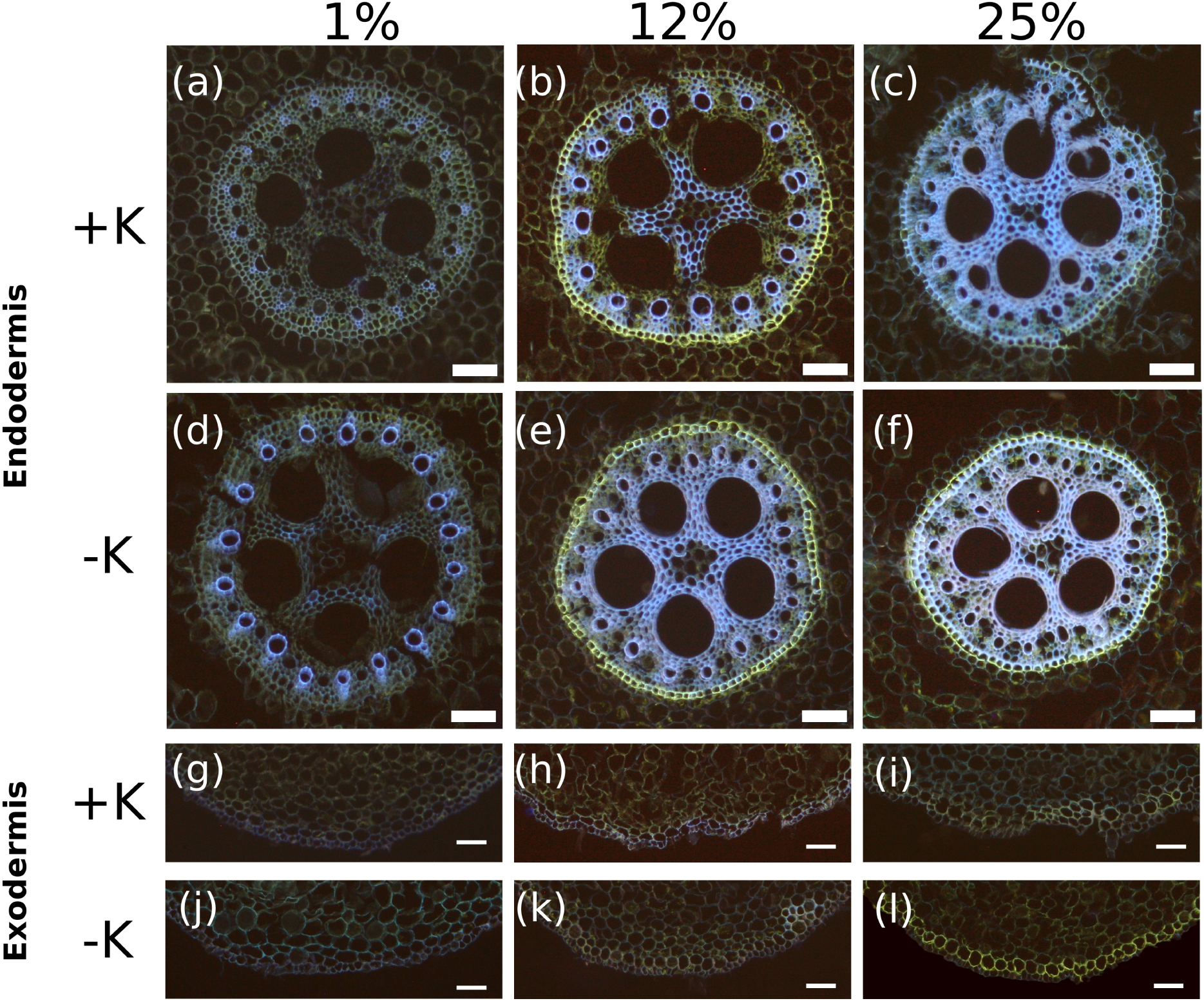
Anatomy of soil-grown maize seminal roots (Fluorol Yellow 088). Fluorol Yellow 088 staining of seminal root cross-sections of maize grown under +K or −K soil conditions, imaged at 1%, 12%, and 25% of total seminal root length from the apex. (a–c) Endodermis under +K and (d–f) endodermis under −K; (g–i) exodermis under +K and (j–l) exodermis under −K. Bright yellow fluorescence indicates suberin lamellae. Scale bars, 100 μm. **Alt text:** Twelve fluorescence micrographs in a grid. Columns are positions at 1, 12 and 25 percent of root length. The upper two rows show endodermis cross-sections under plus K and minus K, the lower two rows exodermis sections. Yellow fluorescence marking suberin lamellae appears earlier and more completely under minus K in the endodermis, while the exodermis differs little between treatments.

Passage cells were present in the endodermis of +K plants but completely absent under K deficiency (Fig. 6a), consistent with full endodermal suberization in −K roots. In hydroponics, the proportion of passage cells was highest under K600, lower under K25 and zero under K100, with K25 and K100 not differing significantly (Fig. 6b).

**Fig. 6.**
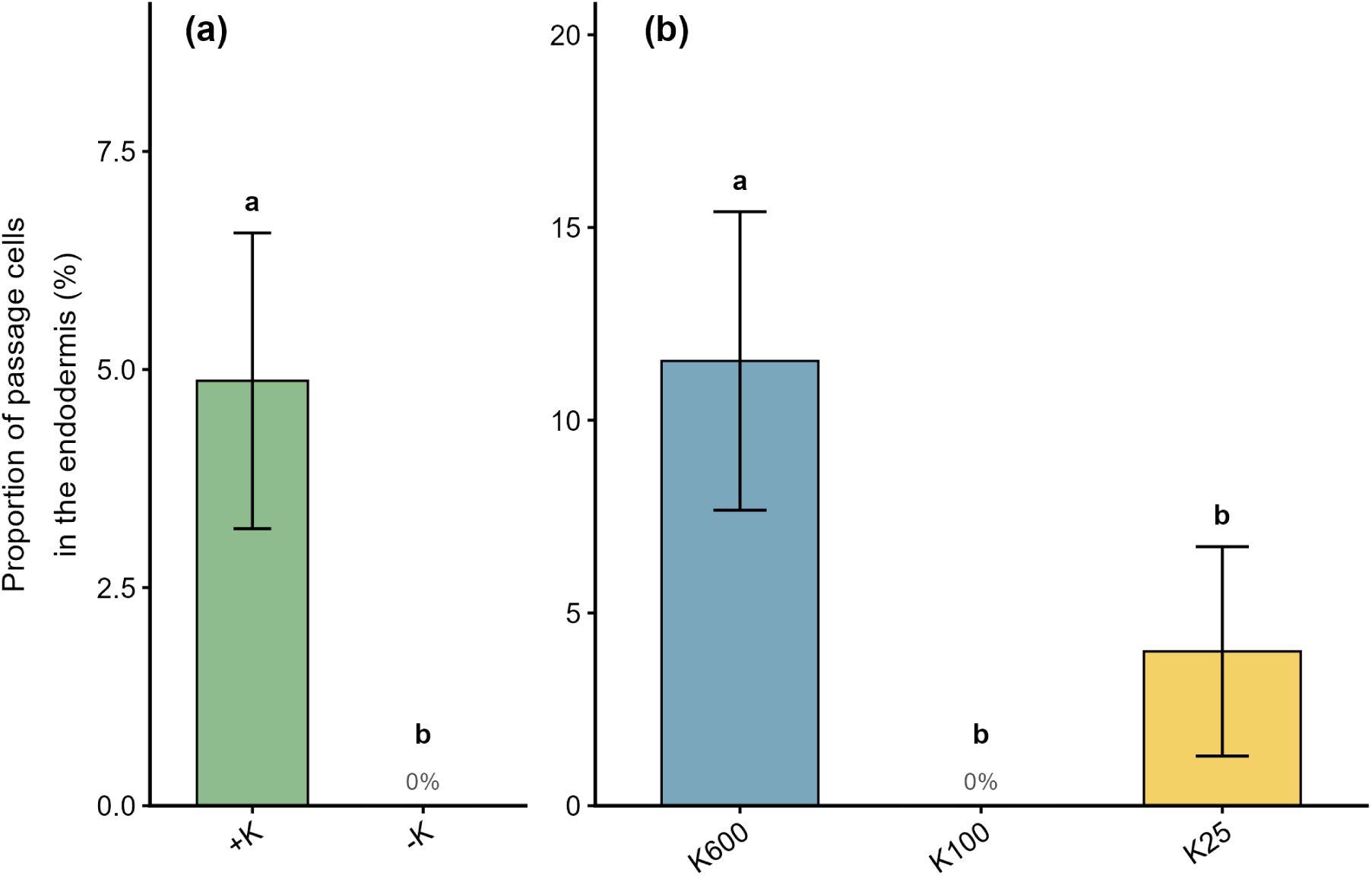
K deficiency eliminates endodermal passage cells. (a) Proportion of passage cells (endodermal cells lacking suberin lamellae) in seminal roots of soil-grown maize under +K and −K conditions. (b) Proportion of passage cells in hydroponically grown plants under the K600, K100, and K25 treatments. Bars represent means ± SD (soil, n = 4; hydroponic, n = 4). Different letters indicate significant differences at α = 0.05 (Tukey’s HSD, two-sided). **Alt text:** Two bar charts of the proportion of endodermal passage cells. Panel a compares soil plants, about 5 percent under plus K and zero under minus K. Panel b compares three hydroponic treatments, about 11 percent under K600, 4 percent under K25 and zero under K100.

Biochemical quantification confirmed that endodermal aliphatic suberin was significantly higher in −K compared to +K plants in Zone A of soil-grown seminal roots, while no significant differences were detected in Zones B or C (Fig. 7; Fig. S2). Exodermal aliphatic suberin did not differ significantly between +K and −K in any zone (Fig. 7), supporting that K deficiency specifically reinforces the endodermal suberized apoplastic barrier.

**Fig. 7.**
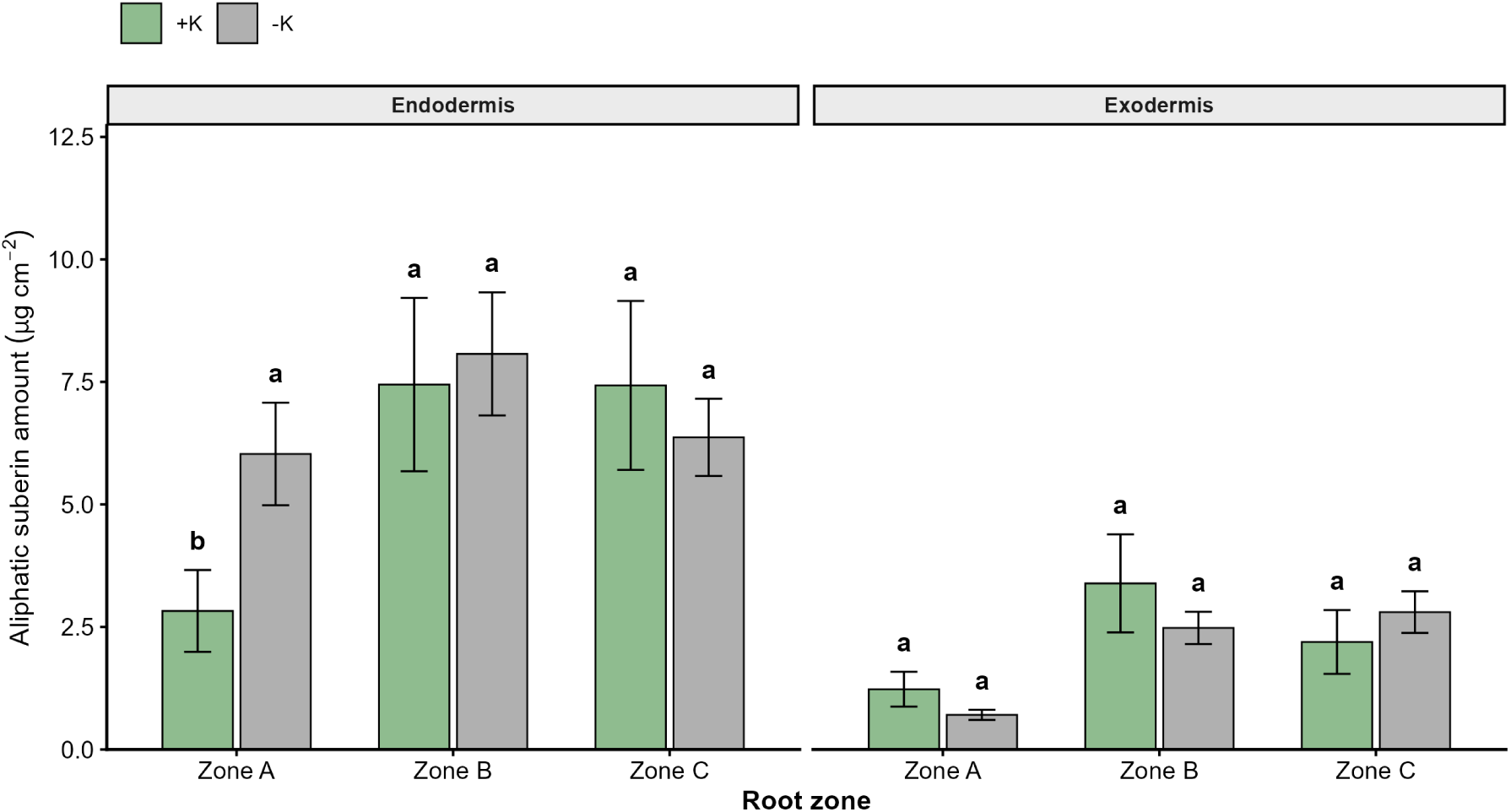
K deficiency selectively increases endodermal aliphatic suberin in soil-grown maize roots. Total aliphatic suberin content in the endodermis (left) and exodermis (right) of seminal roots from maize grown under +K and −K soil conditions, separated into Zone A (0–25%), Zone B (25–50%), and Zone C (50–100% of root length). Bars represent means ± SD (n = 3 biological replicates). Different letters indicate significant differences at α = 0.05 (Tukey’s HSD, two-sided). **Alt text:** Grouped bar chart of aliphatic suberin content in two panels, endodermis on the left and exodermis on the right, each split into root zones A, B and C with plus K and minus K bars side by side. In endodermal zone A the minus K bar is about twice the plus K bar and carries a different letter; all other comparisons share letters.

### Phytohormone treatments modulate endodermal suberin in hydroponically grown roots

Across the hydroponic K gradient, endodermal aliphatic suberin did not differ significantly between K treatments in either root zone, and exodermal aliphatic suberin showed no consistent gradient response (Fig. S9). To test whether ABA contributes to endodermal suberization, hydroponically grown plants were additionally treated with exogenous ABA or with the ABA-biosynthesis inhibitor fluridone. Endodermal aliphatic suberin was higher under both K600+ABA and K600+FLU than under the K600 control in both zones, although this difference was statistically supported only for K600+FLU in Zone A (Fig. S9a,b; Fig. S10). In the exodermis, both hormone treatments also increased aliphatic suberin significantly in Zone B (Fig. S9d). The fluridone results were confounded by severe growth inhibition (see Discussion) and are therefore treated as exploratory. The ABA effect was consistent in direction across both root zones, but remained within the variation of the measurements. The stronger evidence for a contribution of ABA comes from the transcriptional co-regulation of suberin biosynthesis and ABA-responsive genes described below.

### Transcriptomic analysis identifies K deficiency-responsive gene modules associated with endodermal suberin biosynthesis

RNA-seq of K-deficient soil-grown roots identified 848 upregulated and 1192 downregulated genes (|log₂FC| ≥ 1, adjusted p ≤ 0.05; Fig. S3; Table S4). GO enrichment of these genes was dominated by transport-related molecular functions, in particular channel activity and passive transmembrane transporter activity, in which 26 of 27 differentially expressed genes were downregulated (Fig. S4).

WGCNA of hydroponic root transcriptomes identified eight co-expression modules in RNA-seq zone A (10–20% of root length; β = 3) and 15 modules in RNA-seq zone B (40–50%; β = 4), with modules arbitrarily named by colour (Fig. 8). In zone A, the turquoise module (3720 genes) showed the strongest trait associations. It was negatively correlated with endodermal aliphatic suberin (r = −0.87, p = 5×10⁻⁸) and positively correlated with shoot and root dry weight (r = 0.72 and 0.78, respectively). As the sign of a module eigengene is arbitrary in WGCNA, the strength of this association (|r| = 0.87) identifies turquoise as the module most closely linked to endodermal suberin; its positive association with growth and inverse association with deposited suberin reflect the developmental coupling between root growth rate and the position-dependent progression of suberization. In zone B, the blue module was most strongly associated with biomass (root dry weight r = 0.86) and negatively correlated with endodermal aliphatic suberin (r = −0.67; Fig. 8).

**Fig. 8.**
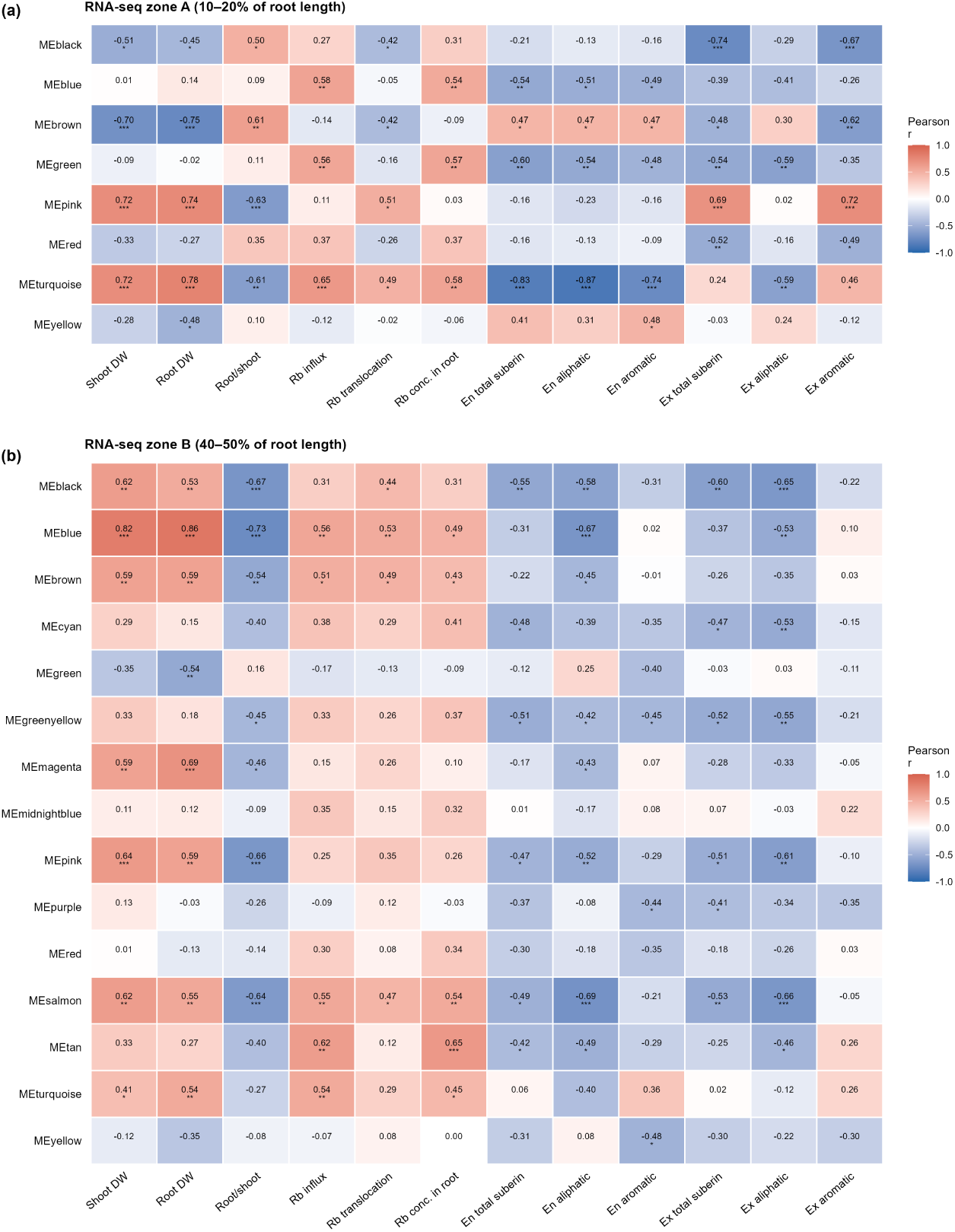
WGCNA co-expression modules associated with endodermal suberin deposition and K deficiency traits. Module-trait correlation heatmap from weighted gene co-expression network analysis (WGCNA) of RNA-seq data from hydroponically grown maize seminal roots in (a) RNA-seq zone A (10–20% of root length; soft-thresholding power β = 3; 8 modules) and (b) RNA-seq zone B (40–50%; β = 4; 15 modules). Co-expression modules are arbitrarily named by colour. Each cell shows the Pearson correlation coefficient and, in parentheses, the corresponding two-sided p-value; asterisks indicate significance (*p < 0.05, **p < 0.01, ***p < 0.001). Traits include shoot dry weight, root dry weight, root-to-shoot ratio, Rb⁺ influx and translocation, root Rb concentration, and endodermal and exodermal total, aliphatic, and aromatic suberin content. The colour scale ranges from blue (negative) to red (positive correlation). **Alt text:** Two module-trait correlation heatmaps from WGCNA, panel a for RNA-seq zone A with eight modules and panel b for zone B with fifteen. Rows are co-expression modules, columns twelve traits from biomass through rubidium fluxes to endodermal and exodermal suberin. Cells are coloured blue for negative and red for positive correlation and carry the coefficient, p value and significance asterisks.

### Conservation and divergence of the transcriptomic K deficiency response between soil and hydroponic systems

To assess whether the transcriptomic K deficiency response is conserved between experimental systems, we compared DEGs from soil-grown roots (2040 DEGs; −K vs. +K) with those from hydroponic RNA-seq zone A (2258 DEGs; K25 vs. K600). Differentially expressed gene numbers across treatments and the overlap between RNA-seq zones A and B are summarized in Fig. S6 and Fig. S7. Only 248 genes were differentially expressed in both datasets (12.2% of soil DEGs, 11.0% of hydroponic DEGs; Fig. S8; Table S3), and of these only 157 were regulated in the same direction in both systems, while 91 responded in opposite directions. Despite this, both systems converged on the same endodermis-specific anatomical response, with loss of endodermal passage cells under K deficiency and no equivalent change in the exodermis. The increase in endodermal aliphatic suberin itself was resolved in the soil experiment.

The suberin biosynthesis genes that emerged as hub genes of the K deficiency-associated co-expression modules (*CYP86A1*, *FAR1*, and *KCS5* homologues; Table S2) were not themselves among this shared set. The convergence on endodermal suberization nonetheless supports a model in which the endodermal suberin response is ABA-dependent, with several K transport genes co-expressed in the same module pointing to coordinated regulation of K transport and apoplastic barrier capacity.

Hub genes of the turquoise module (zone A) and blue module (zone B) included key suberin biosynthesis genes (Table S2): two *CYP86A1* homologues (Zm00001eb367160, Zm00001eb147560) encoding ω-hydroxylases of aliphatic suberin monomer biosynthesis, a *FAR1* fatty acyl-CoA reductase (Zm00001eb201250), and *KCS5* (Zm00001eb190120) mediating very-long-chain fatty acid elongation. A phenylalanine ammonia-lyase (*PAL*; Zm00001eb247660) and a 4-coumarate:CoA ligase (Zm00001eb308860) of the phenylpropanoid pathway were also among the zone A hub genes, and an ABCG-family transporter (Zm00001eb161790) among those of zone B. Additional suberin- and phenylpropanoid-related genes were co-expressed within the same module as non-hub members (kME < 0.80), including a second *FAR1* reductase (Zm00001eb308150), an omega-hydroxypalmitate O-feruloyl transferase (Zm00001eb364060), and cinnamyl-alcohol dehydrogenase *CAD6* (Zm00001eb088800). Several K transport genes were co-expressed within the turquoise module of zone A, including a KT/KUP/HAK-family transporter (Zm00001eb314520; kME 0.68) and the Shaker-type channel *AKT2* (Zm00001eb288580; kME 0.58), both differentially expressed under K deficiency (Table S4). This co-expression of K transport and suberin-barrier genes in the young root zone is consistent with coordinated transcriptional regulation of K transport capacity and apoplastic barrier formation under K deficiency, although the turquoise module is large (3720 genes) and shared module membership alone provides only weak evidence of co-regulation.

## Discussion

The present study demonstrates that K deficiency selectively reinforces endodermal suberized apoplastic barriers in maize seminal roots without equivalent changes in the exodermis. This selectivity was observed consistently across two independent experimental systems, soil-grown and hydroponically grown plants, at the anatomical and transcriptomic levels, while the biochemical increase in endodermal suberin was resolved in the soil experiment. The finding that the exodermal barrier is not reinforced under K deficiency suggests a tissue-specific regulation that may serve to balance K retention with continued access to soil water and nutrients through the outer root layers.

### Endodermal suberization under K deficiency is consistent with a role for ABA signalling

The induction of endodermal suberization in response to K deficiency is consistent with observations in Arabidopsis, where K and sulphur deficiencies were shown to stimulate suberin deposition through endodermal ABA signalling (Barberon et al., 2016). Low K supply is well documented to stimulate ABA production in plants (Wang et al., 2013; Réthoré et al., 2021), which among other functions contributes to K deficiency-induced stomatal closure (Hawkesford et al., 2023). The enhanced endodermal suberization observed here may therefore be associated with increased endogenous ABA production. In contrast, deficiencies of micronutrients such as Fe, Mn, and Zn lead to lower suberin levels via ethylene, highlighting the antagonistic roles of these two hormonal pathways in adapting endodermal permeability to prevailing nutritional conditions (Barberon et al., 2016; Barberon, 2017). In hydroponically grown maize, exogenous ABA tended to increase endodermal aliphatic suberin relative to the K600 control (Fig. S9), consistent with the established role of ABA as a positive regulator of endodermal suberin deposition under abiotic stress (Barberon et al., 2016; Shukla et al., 2021; Meng et al., 2025). Exogenous ABA also increased exodermal suberin in the mature root zone, in line with the requirement of ABA for exodermal suberization in rice (Shiono et al., 2022). ABA is therefore capable of promoting suberization in both layers, and the selectivity observed here reflects the endogenous response to K limitation rather than a restriction of ABA action to the endodermis.

The response to fluridone ran counter to the expectation from an inhibitor of ABA biosynthesis: rather than reducing suberization, it increased endodermal aliphatic suberin in Zone A relative to the K600 control (Fig. S9a). Fluridone inhibits phytoene desaturase and blocks carotenoid biosynthesis upstream of ABA, and at the concentration applied here it was strongly phytotoxic, bleaching the shoots and almost completely arresting growth (Fig. 1, Fig. 2). The dominant effect was therefore unlikely to be a clean reduction of ABA, but rather a generalised carotenoid-depletion stress that is itself a potent inducer of suberin deposition. Fluridone thus does not provide loss-of-function evidence here, and the contribution of ABA rests more securely on the transcriptional co-regulation of suberin-biosynthesis and ABA-responsive genes.

The observed reduction in passage cell proportion to zero under K deficiency in soil-grown plants is particularly striking. Passage cells lack suberin lamellae and therefore remain the sites at which solutes can still be taken up from the apoplast into the symplast across the endodermis. If passage cells are absent, radial transport can continue only for solutes that have already entered the symplast in the outer cortex, from where they move onwards through plasmodesmata. In soil, complete endodermal suberization may carry little cost in older root zones, because the soil solution surrounding them is already depleted by diffusion limitation, so that the uptake capacity provided by unsuberized endodermal cells is of limited value there. A similar elimination of passage cells under osmotic stress was previously reported in barley (Kreszies et al., 2019), suggesting that this response is a conserved strategy in cereals to maximize retention of limiting solutes under stress.

#### Tissue-specific selectivity of the suberization response

The absence of exodermal suberization under K deficiency is particularly informative when considered alongside ABA-mediated suberization responses in other systems and root types. Shiono et al. (2022) showed that ABA is required for exodermal suberization in rice adventitious roots to form a barrier to radial oxygen loss (ROL), a waterlogging response suppressed by fluridone. Subsequently, Shiono & Matsuura (2024) extended this finding to barley adventitious roots, demonstrating that exogenous ABA can artificially induce an ROL barrier with hypodermal suberization in a species that does not normally form such a barrier. By contrast, Grünhofer et al. (2021) reported that in barley seminal roots, exogenous ABA promoted endodermal suberization but did not affect the hypodermis, the outer suberized cell layer beneath the rhizodermis. Our data from maize seminal roots under K deficiency show the same pattern: endodermal suberization is induced while the exodermis remains unresponsive.

These findings reveal a root-type difference in ABA-responsive suberization. Adventitious roots respond to ABA with exodermal suberization, forming a physical barrier against oxygen loss under waterlogging, whereas seminal roots respond with endodermal suberization, reinforcing nutrient retention under deficiency stress. This difference likely reflects tissue-specific expression of ABA signalling components and downstream MYB transcription factors (Shukla et al., 2021), and may represent an evolutionary specialisation matching barrier placement to the primary stress encountered by each root type.

A key finding of the present study is that exodermal suberization was not significantly altered by K deficiency, neither in soil-grown nor in hydroponically grown plants. This is in contrast to the endodermis, which showed clear induction of aliphatic suberin deposition in Zone A under K deficiency. The exodermis of maize is constitutively suberized under optimal conditions (Schreiber et al., 1999), and its aliphatic and aromatic suberin content was already high under the +K control. It is therefore possible that this already high constitutive level limits further induction, or alternatively that the exodermis lacks the requisite ABA-responsive transcriptional machinery for K deficiency-induced upregulation.

The selective endodermal response is biologically advantageous. The exodermis constitutes the outermost suberized barrier and controls radial water and solute influx from the rhizosphere. Under K deficiency, it would be counterproductive to further restrict K entry from the soil by enhancing exodermal suberization. Instead, maintaining exodermal permeability while reinforcing the endodermal barrier serves two functions. It allows continued K uptake from the soil through high-affinity transporters in the epidermis and outer cortex, while preventing K already loaded into the stele from leaking back into the cortical apoplast. This interpretation is supported by the Rb⁺ flux data. K25 plants maintained high Rb⁺ influx capacity, consistent with upregulation of high-affinity K⁺ uptake systems, while root-to-shoot translocation was significantly reduced relative to K600. This suggests preferential K retention in the root symplast under K deficiency.

It is well established that root Na uptake increases under low K supply (Miyamoto et al., 2015; Hawkesford et al., 2023), at least partly because of the antagonistic interaction between these two monovalent cations. Based on the present results, it may also be hypothesized that enhanced Na uptake in K-deficient plants is associated with the absence of additional suberin deposition in the exodermis, which has been described as an outer barrier restricting Na entry (Liu & Kreszies, 2023). Future studies should therefore investigate how K supply affects exodermal suberization in species or genotypes differing in their capacity for Na uptake.

### Reduced root-to-shoot K translocation under K deficiency is consistent with endodermal barrier reinforcement

The Rb⁺ translocation data revealed a striking dissociation between K⁺ influx and root-to-shoot transport under K deficiency. At K25, high-affinity uptake was fully operational, with Rb⁺ influx among the highest across all treatments, yet root-to-shoot translocation was the lowest of all K-gradient treatments. This pattern implies that under sustained K limitation without resupply, K⁺ is preferentially retained in the root rather than being loaded into the xylem for shoot transport. Whether and how rapidly translocation resumes upon K resupply was not addressed in the present experiments. Enhanced endodermal suberization in Zone A, as observed in soil-grown plants, would reduce apoplastic leakage from the stele into the cortex and thereby contribute to maintaining K concentration in the xylem sap. Our results support the hypothesis that endodermal suberin restricts radial K⁺ leakage from the stele. Using caesium as a K⁺ tracer and laser-ablation ICP-MS imaging, Vestenaa et al. (2024) found that more heavily suberized root zones showed reduced tracer leakage and correspondingly higher K⁺ translocation efficiency in both Arabidopsis mutants and barley seminal roots. A complementary line of evidence comes from CRISPR/Cas9-induced suberin-deficient poplar roots, in which loss of suberization increased the translocation of NaCl and of a photosynthesis-inhibiting herbicide, consistent with a reduced solute barrier function of endodermal suberin lamellae (Grünhofer et al., 2024). Shoot K concentrations were not significantly altered in suberin-deficient mutants across multiple Arabidopsis studies (Grünhofer et al., 2024, their Table 1). This suggests that K homeostasis can be maintained by active transport systems even when the apoplastic barrier is compromised. Restriction of passive leakage from the stele therefore remains the more physiologically relevant function of endodermal suberization under K deficiency.

Fluridone also reduced Rb⁺ influx and translocation to very low values, but this coincided with near-complete growth arrest (Fig. 2) and therefore cannot be ascribed specifically to a loss of ABA signalling rather than to generalised stress. Independently of the fluridone treatment, several K transport genes were co-expressed with core suberin biosynthesis genes in the turquoise module of RNA-seq zone A. This suggests that K transport and endodermal suberization are transcriptionally co-regulated in maize roots under K deficiency, a relationship that warrants further functional characterization.

### Transcriptomic co-regulation of K⁺ transport and suberin biosynthesis

The WGCNA analysis identified the turquoise module in RNA-seq zone A (10–20% of root length) as the primary K deficiency-responsive co-expression module, negatively correlated with endodermal aliphatic suberin (r = −0.87) and positively with shoot dry weight. Hub genes within this module included *CYP86A1* homologues and a *FAR1* fatty acyl-CoA reductase, both encoding enzymes of the aliphatic suberin biosynthesis pathway (Table S2). *KCS5* was a hub gene of the zone B module rather than of turquoise. Their steady-state transcript levels did not rise with K deficiency, consistent with the negative correlation between the module eigengene and deposited suberin and with the reduced expression of canonical biosynthesis genes (e.g. *KCS5*) in K-deficient roots. As deposited suberin is a cumulative product that integrates earlier biosynthetic activity, increased deposition need not coincide with elevated transcript levels at the sampled position; rather, the programme is ABA-responsive (in zone A induced by exogenous ABA and repressed by fluridone; Fig. S5), with lamellae continuing to accumulate from enzymes expressed earlier in development, apical to the sampled zone. These genes are maize orthologues of well-characterized Arabidopsis suberin biosynthesis genes (Franke & Schreiber, 2007; Pollard et al., 2008; Compagnon et al., 2009). Their co-expression with genes of the general phenylpropanoid pathway, including *PAL*, 4-coumarate:CoA ligase, laccases and peroxidases, indicates coordinated regulation of aliphatic suberin biosynthesis and phenylpropanoid metabolism. As these enzymes supply precursors for both the aromatic domain of suberin and for lignin, this co-expression cannot be attributed to suberin specifically; endodermal differentiation involves lignification as well. An ABCG-family transporter was a hub gene of the zone B module; ABCG proteins include the known exporters of suberin monomers in Arabidopsis, although the function of this maize homologue has not been tested. In contrast to the tight co-expression of suberin biosynthesis genes within the turquoise module of zone A, the corresponding genes were distributed across several modules in zone B. This dispersal is consistent with a relaxation of the co-regulated suberization programme in the older, largely constitutively suberized root zone. It supports the view that K deficiency-induced endodermal suberization is concentrated in the younger, actively differentiating zone A.

Two homologues of the transcription factor *MYB36* (Zm00001eb086570, kME 0.94; Zm00001eb313580, kME 0.81) were hub genes of the zone B module. In Arabidopsis, *MYB36* is the master regulator of Casparian strip formation rather than of suberin lamellae (Kamiya et al., 2015), so their appearance here points to coordinated control of endodermal differentiation more broadly. Suberin lamellae themselves are controlled by a separate set of MYB factors, *MYB41*, *MYB53*, *MYB92* and *MYB93* (Shukla et al., 2021), with analogous MYB-dependent regulation described in barley under aluminium stress (Meng et al., 2025); no clear homologue of these was among the hub genes of either module. The maize *MYB36* homologues nevertheless provide candidate regulators for future functional studies in a crop species of global agricultural importance.

A recent transcriptomic study of maize roots under K deficiency identified 5972 DEGs and revealed enrichment of K⁺ signalling and transporter-related genes (Guo et al., 2023). Our study expands on these findings by linking transcriptomic responses specifically to the onset of suberization in defined root zones and by providing biochemical quantification of the resulting barrier changes. The moderate overlap of 12% between the soil and hydroponic DEG datasets highlights the complexity of the K deficiency response across the two systems.

A methodological consideration applies to these module-trait correlations. Whereas suberin was quantified separately for endodermis and exodermis by enzymatic isolation of the respective cell wall layers, RNA was extracted from whole root segments, in which the endodermis represents only a small fraction of the cells. The transcriptomic signal is therefore tissue-averaged: genes assigned here to the suberization response may in part be expressed in adjacent cell types, and a genuinely endodermal signal may be diluted by the surrounding tissues. Spatially resolved approaches, such as laser capture microdissection of the endodermis (Meng et al., 2025) or single-cell RNA-sequencing, will be required to assign these programmes unambiguously.

### Convergent physiological outcomes despite divergent transcriptomes in soil and hydroponic systems

The comparison of K deficiency-responsive transcriptomes between soil-grown and hydroponically grown plants revealed a relatively modest overlap of 248 genes (12.2% of soil DEGs, 11.0% of hydroponic DEGs). The complexity of soil signals, including microbiome interactions, organic matter, and pH, generates a substantially broader transcriptomic response than defined hydroponic solutions (Kreszies et al., 2019; Guo et al., 2023). A further difference concerns the K concentration that roots actually experience. A K-deficient nutrient solution is uniformly deficient along the entire root system, whereas in soil the concentration at the root surface is set locally by diffusion, so that the solution immediately around a growing root becomes depleted even when the bulk soil is adequately supplied, most strongly in older root zones. Roots in soil therefore experience a spatial and temporal gradient of K availability that has no counterpart in hydroponics, and this more gradual depletion may buffer the transcriptomic response relative to the acute contrast imposed by K25 versus K600.

Despite this transcriptomic divergence, both experimental systems produced the same endodermis-specific anatomical response, without equivalent changes in the exodermis; the biochemical increase in endodermal suberin was resolved in the soil experiment. This convergence suggests that the endodermal suberization response is a robust and conserved adaptive mechanism activated across a wide range of K deficiency conditions. Although the principal suberin biosynthesis genes (*CYP86A1*, *FAR1*, and *KCS5* homologues) were not themselves among the shared set, the 157 genes regulated in the same direction in both systems may represent a core K deficiency-responsive programme. A comparison with nitrogen deficiency in the same cultivar is informative (Liu et al., 2026). Under early nitrogen limitation, maize seminal roots likewise reinforced the endodermis rather than the exodermis. However, the suberin response was comparatively weak and restricted to the aromatic fraction, with no significant change in aliphatic suberin, and the dominant anatomical adjustment was instead an increase in aerenchyma formation. K deficiency, by contrast, elicited a clear increase in endodermal aliphatic suberin. Both nutrients therefore act selectively on the endodermis rather than the exodermis, yet the nature and strength of the response differ. This indicates that nutrient-induced endodermal plasticity is tuned not only to the species and tissue but also to the specific nutrient and to the metabolic cost of the adaptation.

### Implications for K use efficiency in maize

Maize is among the most K-demanding crop species and is frequently grown on K-depleted soils, particularly in developing regions where fertiliser inputs are constrained. The present study demonstrates that maize roots activate an adaptive suberization response under K deficiency. This response appears functionally analogous to the nutrient-induced endodermal plasticity described in Arabidopsis (Barberon et al., 2016), but with tissue-specific differences unique to the maize root anatomy, which harbours both a constitutively suberized exodermis and a developmentally regulated endodermis. Understanding the molecular control of this tissue-specific response may open avenues for engineering crops with optimized K retention capacity under low-input conditions. The lack of an exodermal response is itself consistent with a functional division of labour: reinforcing the outer barrier would restrict K entry without further limiting loss from the stele, whereas reinforcing the endodermis restricts loss while leaving uptake capacity intact. The constitutively suberized exodermis is nevertheless likely to serve functions beyond K nutrition that were not addressed here; in potato, reduced exodermal suberin alters element concentrations and impairs growth (Company-Arumí et al., 2026).

The co-regulation of K transport genes with endodermal suberization in the same transcriptional module raises the intriguing possibility that K⁺ uptake capacity and apoplastic barrier function are part of a single adaptive programme, potentially coordinated upstream by ABA signalling. Future studies combining promoter analyses of the hub genes with targeted genetic validation, for example CRISPR/Cas9 editing of candidate K transporters (such as Zm00001eb314520, KT/KUP/HAK family) or of the candidate MYB regulators, would be valuable in dissecting this regulatory architecture. Together, our data establish maize as a tractable system for investigating the physiological and molecular mechanisms of nutrient-induced apoplastic barrier plasticity in a crop species context.

## Acknowledgements

We thank Kirsten Fladung and Ulrike Kierbaum for technical assistance, Hendrik Könning for taking the plant photographs, and Alexander Geuer and Heinrich Georg Meyer for support with the hydroponic experiments. We are grateful to the Landesamt für Landwirtschaft und Ländlichen Raum Thüringen, Dr Günter Kießling, and Dr Wilfried Zorn for providing the original low-K soil.

## Funding

This work was supported by the China Scholarship Council (CSC) through a doctoral scholarship to TL and by K+S Minerals and Agriculture GmbH.

## Conflict of interest

No conflict of interest declared.

## Author contributions

Conceptualization: TK, TL, KD, IC, JG, LS. Investigation: TL, PG, VZD, LS. Formal analysis: TL, TK. Resources: KD, IC, JG, LS. Writing – original draft: TL, TK. Writing – review & editing: all authors. All authors read and approved the final manuscript.

## Data availability

The RNA-sequencing raw reads generated in this study are available in the NCBI Sequence Read Archive under BioProject accession PRJNA1517815. All other data supporting the findings of this study are available within the article and its Supporting Information, or from the corresponding author on reasonable request.

## Acknowledgement of AI use

During the preparation of this work the authors used an AI-based assistant (Claude, Anthropic) for language editing, for restructuring parts of the text, and for cross-checking data, and figures against the underlying source files. The assistant was not used to generate data, to design the experiments, or to produce the primary analyses. All content was reviewed and verified by the authors, who take full responsibility for the work.

## Abbreviations

ABA: abscisic acid
DEG: differentially expressed gene
DW: dry weight
FLU: fluridone
FPKM: fragments per kilobase of transcript per million mapped reads
FY: Fluorol Yellow 088
GO: Gene Ontology
GS: gene significance
KEGG: Kyoto Encyclopedia of Genes and Genomes
kME: module membership
WGCNA: weighted gene co-expression network analysis.

## Supporting Information

The following Supporting Information is available for this article.

**Fig. S1 Soil moisture during the cultivation of maize.** Soil moisture (% of the maximum water-holding capacity) recorded throughout cultivation by sensors placed in the upper (top 1–6) and lower (bottom 1–6) soil layers.

**Alt text:** Line chart of soil moisture over the cultivation period, with twelve traces for sensors in the upper and lower soil layer. The vertical axis runs from 40 to 100 percent of the maximum water-holding capacity. All traces fluctuate between roughly 65 and 87 percent and drift slightly downward over time.

**Fig. S2 Aliphatic suberin substance classes in the endodermis and exodermis of soil-grown maize seminal roots.** (a) Endodermis; (b) exodermis, each shown for Zones A–C. Substance classes are primary alcohols (alc), unsubstituted fatty acids (fa), α,ω-dicarboxylic acids (diacids), ω-hydroxy acids (ω-OH) and 2-hydroxy acids (2-OH). In the endodermis of Zone A, 2-hydroxy acids were close to the detection limit under +K, so the corresponding bar is not visible. Bars represent means ± SD (n = 3 biological replicates).

**Alt text:** Bar charts of five aliphatic suberin substance classes, panel a endodermis and panel b exodermis, each shown for root zones A to C with plus K and minus K bars side by side. The classes are primary alcohols, unsubstituted fatty acids, dicarboxylic acids, omega-hydroxy acids and 2-hydroxy acids. Bars are means with standard deviation.

**Fig. S3 Differentially expressed genes under K deficiency in soil-grown maize roots.** Volcano plot of log₂ fold change (−K vs +K) against statistical significance. Genes with | log₂FC| ≥ 1 and adjusted p ≤ 0.05 were considered differentially expressed (two-sided Wald test, Benjamini–Hochberg correction) (848 upregulated, 1192 downregulated).

**Alt text:** Volcano plot of the soil transcriptome. The horizontal axis is the log2 fold change of minus K against plus K, the vertical axis the negative logarithm of the adjusted p value. Downregulated genes are blue on the left and upregulated genes red on the right, with dashed threshold lines and counts of 848 upregulated and 1192 downregulated genes. The twenty most extreme genes are labelled.

**Fig. S4 Gene Ontology enrichment among K deficiency-responsive genes in soil-grown maize roots.** Molecular function terms significantly enriched (one-sided hypergeometric test, adjusted p ≤ 0.05, Benjamini–Hochberg) among the 2040 differentially expressed genes shown in Fig. S3, tested against all 13,805 GO-annotated genes of the soil dataset. Bubble size gives the number of differentially expressed genes annotated with the term, and the dashed line marks adjusted p = 0.05. These five terms are the complete set of significantly enriched terms; no biological process or cellular component term reached significance.

**Alt text:** Bubble plot of five enriched molecular function terms, from most to least significant: channel activity, passive transmembrane transporter activity, ADP binding, O-methyltransferase activity and glucosyltransferase activity. The horizontal axis is the negative logarithm of the adjusted p value with a dashed line at p equals 0.05, and bubble size gives the number of differentially expressed genes annotated with each term.

**Fig. S5 Expression of suberin biosynthesis and phenylpropanoid genes in hydroponic maize root zones A and B.** (a) RNA-seq zone A (10–20% of root length); (b) RNA-seq zone B (40–50%). Shown are genes differentially expressed in at least two treatments relative to the K600 control (adjusted p ≤ 0.05, no fold-change threshold, and therefore a wider set than the differentially expressed genes counted elsewhere in this study); colour indicates log₂ fold change relative to K600. The K100 treatment is not shown in (a) because none of these genes was differentially expressed under K100 in zone A. Rows are grouped by pathway (aliphatic suberin biosynthesis, phenylpropanoid) as indicated by the annotation bar.

**Alt text:** Two heatmaps of gene expression relative to the K600 control, panel a for zone A with sixteen genes and panel b for zone B with twelve. Rows are individual suberin biosynthesis and phenylpropanoid genes with identifier and annotation, columns the remaining treatments. Colour runs from blue for lower to red for higher expression, and a side bar marks the pathway of each gene.

**Fig. S6 Numbers of differentially expressed genes across treatments in hydroponically grown maize roots.** (a) RNA-seq zone A (10–20% of root length); (b) RNA-seq zone B (40–50%). Bars give the genes differentially expressed relative to the K600 control (|log₂FC| ≥ 1, adjusted p ≤ 0.05), with upregulated genes plotted above and downregulated genes below the zero line; the numbers next to the bars are the gene counts. Note that the y-axis scales differ between (a) and (b).

**Alt text:** Two diverging bar charts of differentially expressed gene numbers relative to the K600 control, panel a for zone A and panel b for zone B. Upregulated genes are plotted in red above the zero line and downregulated genes in blue below it, with counts printed next to each bar. Fluridone produces by far the largest response in both zones.

**Fig. S7 Overlap of K deficiency-responsive genes between RNA-seq zone A (10–20% of root length) and zone B (40–50%) of hydroponically grown maize roots (K25 vs K600).** Venn diagram of the genes differentially expressed between K25 and K600 (|log₂FC| ≥ 1, adjusted p ≤ 0.05) in each zone: 2168 in zone A only, 114 in zone B only and 90 in both.

**Alt text:** Venn diagram with two overlapping circles for the K25 against K600 comparison. The zone A circle holds 2168 genes alone, the zone B circle 114 alone, and 90 genes lie in the overlap.

**Fig. S8 Comparison of K deficiency-responsive transcriptomes between soil and hydroponic experiments.** Overlap between differentially expressed genes of soil-grown roots (−K vs +K; 2040 genes) and of hydroponic RNA-seq zone A (K25 vs K600; 2258 genes); the 248 shared genes are listed in Table S3.

**Alt text:** Panel a is a Venn diagram of the soil and hydroponic zone A gene sets, with 1792 genes only in soil, 2010 only in hydroponics and 248 shared. Panel b is a horizontal bar chart splitting these into seven categories: 59 shared and concordantly upregulated, 98 shared and concordantly downregulated, 91 shared but opposite in direction, and the soil-only and hydroponics-only genes by direction.

**Fig. S9 Endodermal and exodermal aliphatic suberin in hydroponically grown maize seminal roots under K supply and phytohormone treatments.** (a, b) Endodermis in Zone A (0–25% of root length) and Zone B (25–50%); (c, d) exodermis in the same zones. Bars represent means ± SD (n = 3, except endodermis under K600+ABA and K600+FLU in Zone B, where n = 2). Lowercase letters indicate significant differences among K-gradient treatments, uppercase letters among hormone-modified treatments (Tukey’s HSD, two-sided, α = 0.05).

**Alt text:** Four bar charts of aliphatic suberin in hydroponically grown roots. Panels a and b show the endodermis in zones A and B, panels c and d the exodermis in the same zones, each across the six treatments. Bars are means with standard deviation and carry lowercase letters for the K gradient and uppercase letters for the hormone treatments.

**Fig. S10 Anatomy of hydroponically grown maize seminal roots across K supply and phytohormone treatments.** Cross-sections stained with Fluorol Yellow 088 and imaged under UV excitation; suberin lamellae appear as yellow fluorescence in the endodermis and exodermis. Columns give the position along the seminal root as a percentage of total root length (25%, 50%, 70%); rows give the treatment. (a–c) K25, (d–f) K100, (g–i) K600, (j–l) K5000, (m–o) K600+ABA and (p–r) K600+FLU. Scale bars, 100 μm.

**Alt text:** Eighteen fluorescence micrographs of root cross-sections in a grid of three columns and six rows. Columns are positions at 25, 50 and 70 percent of root length, rows the treatments K25, K100, K600, K5000, K600 plus ABA and K600 plus fluridone. Green and yellow fluorescence marks suberized endodermal and exodermal cell walls.

**Table S1 Soil properties of the substrate used for the soil experiment.** Texture, pH, and nutrient status of the soil determined prior to the experiment.

**Table S2 Hub genes of WGCNA modules associated with suberin biosynthesis and potassium transport.** Hub genes were defined as module members with module membership kME ≥ 0.80 and absolute gene significance for endodermal aliphatic suberin > 0.5, listed separately for the turquoise module of RNA-seq zone A and the blue module of zone B. A separate worksheet lists all co-expression modules of both zones with their gene numbers; the module assignment of every gene is given in Table S4.

**Table S3 Genes differentially expressed under K deficiency in both soil-grown and hydroponic maize roots**. The 248 genes differentially expressed under K deficiency in both systems, with their fold changes in each; 157 respond in the same direction and 91 in opposite directions.

**Table S4 Differentially expressed genes across treatment comparisons in soil-grown and hydroponic maize roots**. Differentially expressed genes for each treatment comparison (adjusted p ≤ 0.05 and |log₂FC| ≥ 1), with fold changes, expression means, and WGCNA module membership of the matched RNA-seq zone. One worksheet per comparison plus an overview sheet.

## Notes

### Competing Interest Statement

The authors have declared no competing interest.

## References

Baales J, Zeisler-Diehl VV, Schreiber L. 2021. Analysis of extracellular cell wall lipids: wax, cutin, and suberin in leaves, roots, fruits, and seeds. In: Bartels D, Dörmann P, eds. Plant lipids. Methods and protocols. New York, NY, USA: Springer US; Humana, 275–293. doi: 10.1007/978-1-0716-1362-7_15

Barberon M. 2017. The endodermis as a checkpoint for nutrients. New Phytologist 213: 1604–1610. doi: 10.1111/nph.14140

Barberon M, Vermeer JEM, De Bellis D, Wang P, Naseer S, Andersen TG, Humbel BM, Nawrath C, Takano J, Salt DE, Geldner N. 2016. Adaptation of root function by nutrient-induced plasticity of endodermal differentiation. Cell 164: 447–459. doi: 10.1016/j.cell.2015.12.021

Beier S, Marella NC, Yvin J-C, Hosseini SA, von Wirén N. 2022. Silicon mitigates potassium deficiency by enhanced remobilization and modulated potassium transporter regulation. Environmental and Experimental Botany 198: 104849. doi: 10.1016/j.envexpbot.2022.104849

Brundrett MC, Kendrick B, Peterson CA. 1991. Efficient lipid staining in plant material with Sudan Red 7B or Fluorol Yellow 088 in polyethylene glycol–glycerol. Biotechnic & Histochemistry 66: 111–116. doi: 10.3109/10520299109110562

Cakmak I. 2005. The role of potassium in alleviating detrimental effects of abiotic stresses in plants. Journal of Plant Nutrition and Soil Science 168: 521–530. doi: 10.1002/jpln.200420485

Chen S, Zhou Y, Chen Y, Gu J. 2018. fastp: an ultra-fast all-in-one FASTQ preprocessor. Bioinformatics 34: i884–i890. doi: 10.1093/bioinformatics/bty560

Compagnon V, Diehl P, Benveniste I, Meyer D, Schaller H, Schreiber L, Franke R, Pinot F. 2009. CYP86B1 is required for very long chain ω-hydroxyacid and α,ω-dicarboxylic acid synthesis in root and seed suberin polyester. Plant Physiology 150: 1831–1843. doi: 10.1104/pp.109.141408

Company-Arumí D, Montells C, Iglesias M, Marguí E, Verdaguer D, Vogel-Mikuš K, Kelemen M, Figueras M, Anticó E, Serra O. 2026. Exodermal suberin contributes to ion homeostasis and growth in potato. Journal of Experimental Botany: erag205. doi: 10.1093/jxb/erag205

Coskun D, Britto DT, Kronzucker HJ. 2014. The physiology of channel-mediated K+ acquisition in roots of higher plants. Physiologia Plantarum 151: 305–312. doi: 10.1111/ppl.12174

Franke R, Schreiber L. 2007. Suberin – a biopolyester forming apoplastic plant interfaces. Current Opinion in Plant Biology 10: 252–259. doi: 10.1016/j.pbi.2007.04.004

Gamble PE, Mullet JE. 1986. Inhibition of carotenoid accumulation and abscisic acid biosynthesis in fluridone-treated dark-grown barley. European Journal of Biochemistry 160: 117–121. doi: 10.1111/j.1432-1033.1986.tb09949.x

Geldner N. 2013. The endodermis. Annual Review of Plant Biology 64: 531–558. doi: 10.1146/annurev-arplant-050312-120050

Grünhofer P, Schreiber L, Kreszies T. 2021. Suberin in monocotyledonous crop plants: structure and function in response to abiotic stresses. In: Baluška F, Mukherjee S, eds. Rhizobiology: molecular physiology of plant roots. New York, NY, USA: Springer, 333–378. doi: 10.1007/978-3-030-84985-6_19

Grünhofer P, Heimerich I, Pohl S, Oertel M, Meng H, Zi L, Lucignano K, Bokhari SNH, Guo Y, Li R, Lin J, Fladung M, Kreszies T, Stöcker T, Schoof H, Schreiber L. 2024. Suberin deficiency and its effect on the transport physiology of young poplar roots. New Phytologist 242: 137–153. doi: 10.1111/nph.19588

Guo S, Liu Z, Sheng H, Olukayode T, Zhou Z, Liu Y, Wang M, He M, Kochian L, Qin Y. 2023. Dynamic transcriptome analysis unravels key regulatory genes of maize root growth and development in response to potassium deficiency. Planta 258: 99. doi: 10.1007/s00425-023-04260-7

Hawkesford MJ, Cakmak I, Coskun D, De Kok LJ, Lambers H, Schjoerring JK, White PJ. 2023. Functions of macronutrients. In: Rengel Z, Cakmak I, White PJ, eds. Marschner’s mineral nutrition of plants, 4th edn. London, UK: Elsevier, 201–281. doi: 10.1016/B978-0-12-819773-8.00019-8

Hose E, Clarkson DT, Steudle E, Schreiber L, Hartung W. 2001. The exodermis: a variable apoplastic barrier. Journal of Experimental Botany 52: 2245–2264. doi: 10.1093/jexbot/52.365.2245

Kamiya T, Borghi M, Wang P, Danku JMC, Kalmbach L, Hosmani PS, Naseer S, Fujiwara T, Geldner N, Salt DE. 2015. The MYB36 transcription factor orchestrates Casparian strip formation. Proceedings of the National Academy of Sciences, USA 112: 10533–10538. doi: 10.1073/pnas.1507691112

Kim D, Paggi JM, Park C, Bennett C, Salzberg SL. 2019. Graph-based genome alignment and genotyping with HISAT2 and HISAT-genotype. Nature Biotechnology 37: 907–915. doi: 10.1038/s41587-019-0201-4

Kreszies T, Schreiber L, Ranathunge K. 2018. Suberized transport barriers in Arabidopsis, barley and rice roots: from the model plant to crop species. Journal of Plant Physiology 227: 75–83. doi: 10.1016/j.jplph.2018.02.002

Kreszies T, Shellakkutti N, Osthoff A, Yu P, Baldauf JA, Zeisler-Diehl VV, Ranathunge K, Hochholdinger F, Schreiber L. 2019. Osmotic stress enhances suberization of apoplastic barriers in barley seminal roots: analysis of chemical, transcriptomic and physiological responses. New Phytologist 221: 180–194. doi: 10.1111/nph.15351

Langfelder P, Horvath S. 2008. WGCNA: an R package for weighted correlation network analysis. BMC Bioinformatics 9: 559. doi: 10.1186/1471-2105-9-559

Liu T, Kreszies T. 2023. The exodermis: a forgotten but promising apoplastic barrier. Journal of Plant Physiology 290: 154118. doi: 10.1016/j.jplph.2023.154118

Liu T, Palisaar J, Grünhofer P, Zeisler-Diehl VV, Schreiber L, Dittert K, Kreszies T. 2026. Nitrogen availability affects aerenchyma formation and suberization in early root development of soil-grown maize. Plant Science 362: 112786. doi: 10.1016/j.plantsci.2025.112786

Love MI, Huber W, Anders S. 2014. Moderated estimation of fold change and dispersion for RNA-seq data with DESeq2. Genome Biology 15: 550. doi: 10.1186/s13059-014-0550-8

Lyzenga WJ, Liu Z, Olukayode T, Zhao Y, Kochian LV, Ham B-K. 2023. Getting to the roots of N, P, and K uptake. Journal of Experimental Botany 74: 1784–1805. doi: 10.1093/jxb/erad035

Meng H, Zhang Q, Kreszies T, Acosta IF, Schreiber L. 2025. Regulatory programmes driving suberin plasticity under aluminium stress in barley roots. Plant, Cell & Environment 48: e70075. doi: 10.1111/pce.70075

Miyamoto T, Ochiai K, Nonoue Y, Matsubara K, Yano M, Matoh T. 2015. Expression level of the sodium transporter gene OsHKT2;1 determines sodium accumulation of rice cultivars under potassium-deficient conditions. Soil Science and Plant Nutrition 61: 481–492. doi: 10.1080/00380768.2015.1005539

Nieves-Cordones M, Alemán F, Martínez V, Rubio F. 2014. K+ uptake in plant roots. The systems involved, their regulation and parallels in other organisms. Journal of Plant Physiology 171: 688–695. doi: 10.1016/j.jplph.2013.09.021

Pertea M, Pertea GM, Antonescu CM, Chang T-C, Mendell JT, Salzberg SL. 2015. StringTie enables improved reconstruction of a transcriptome from RNA-seq reads. Nature Biotechnology 33: 290–295. doi: 10.1038/nbt.3122

Peterson CA, Murrmann M, Steudle E. 1993. Location of the major barriers to water and ion movement in young roots of Zea mays L. Planta 190: 127–136. doi: 10.1007/BF00195684

Pollard M, Beisson F, Li Y, Ohlrogge JB. 2008. Building lipid barriers: biosynthesis of cutin and suberin. Trends in Plant Science 13: 236–246. doi: 10.1016/j.tplants.2008.03.003

Ranathunge K, Lin J, Steudle E, Schreiber L. 2011. Stagnant deoxygenated growth enhances root suberization and lignification, but differentially affects water and NaCl permeabilities in rice (Oryza sativa L.) roots. Plant, Cell & Environment 34: 1223–1240. doi: 10.1111/j.1365-3040.2011.02318.x

Ranathunge K, Kim YX, Wassmann F, Kreszies T, Zeisler V, Schreiber L. 2017. The composite water and solute transport of barley (Hordeum vulgare) roots: effect of suberized barriers. Annals of Botany 119: 629–643. doi: 10.1093/aob/mcw252

Réthoré E, Jing L, Ali N, Yvin J-C, Pluchon S, Hosseini SA. 2021. K deprivation modulates the primary metabolites and increases putrescine concentration in Brassica napus. Frontiers in Plant Science 12: 681895. doi: 10.3389/fpls.2021.681895

Robe K, Barberon M. 2023. Nutrient carriers at the heart of plant nutrition and sensing. Current Opinion in Plant Biology 74: 102376. doi: 10.1016/j.pbi.2023.102376

Sardans J, Peñuelas J. 2021. Potassium control of plant functions: ecological and agricultural implications. Plants 10: 419. doi: 10.3390/plants10020419

Schreiber L, Hartmann K, Skrabs M, Zeier J. 1999. Apoplastic barriers in roots: chemical composition of endodermal and hypodermal cell walls. Journal of Experimental Botany 50: 1267–1280. doi: 10.1093/jxb/50.337.1267

Schüller H. 1969. Die CAL-Methode, eine neue Methode zur Bestimmung des pflanzenverfügbaren Phosphates in Böden. Zeitschrift für Pflanzenernährung und Bodenkunde 123: 48–63. doi: 10.1002/jpln.19691230106

Shiono K, Yoshikawa M, Kreszies T, Yamada S, Hojo Y, Matsuura T, Mori IC, Schreiber L, Yoshioka T. 2022. Abscisic acid is required for exodermal suberization to form a barrier to radial oxygen loss in the adventitious roots of rice (Oryza sativa). New Phytologist 233: 655–669. doi: 10.1111/nph.17751

Shiono K, Matsuura H. 2024. Exogenous abscisic acid induces the formation of a suberized barrier to radial oxygen loss in adventitious roots of barley (Hordeum vulgare). Annals of Botany 133: 931–940. doi: 10.1093/aob/mcae010

Shukla V, Han J-P, Cléard F, Lefebvre-Legendre L, Gully K, Flis P, Berhin A, Andersen TG, Salt DE, Nawrath C, Barberon M. 2021. Suberin plasticity to developmental and exogenous cues is regulated by a set of MYB transcription factors. Proceedings of the National Academy of Sciences, USA 118: e2101730118. doi: 10.1073/pnas.2101730118

Sijmons PC, Kolattukudy PE, Bienfait FH. 1985. Iron deficiency decreases suberization in bean roots through a decrease in suberin-specific peroxidase activity. Plant Physiology 78: 115–120. doi: 10.1104/pp.78.1.115

Steudle E, Peterson CA. 1998. How does water get through roots? Journal of Experimental Botany 49: 775–788. doi: 10.1093/jxb/49.322.775

VDLUFA. 1998. Standpunkt: Kalium-Düngung. Darmstadt, Germany: Verband Deutscher Landwirtschaftlicher Untersuchungs-und Forschungsanstalten

Vestenaa MW, Husted S, Minutello F, Persson DP. 2024. Endodermal suberin restricts root leakage of cesium: a suitable tracer for potassium. Physiologia Plantarum 176: e14393. doi: 10.1111/ppl.14393

Wang M, Zheng Q, Shen Q, Guo S. 2013. The critical role of potassium in plant stress response. International Journal of Molecular Sciences 14: 7370–7390. doi: 10.3390/ijms14047370

Yu G, Wang L-G, Han Y, He Q-Y. 2012. clusterProfiler: an R package for comparing biological themes among gene clusters. OMICS 16: 284–287. doi: 10.1089/omi.2011.0118

Zhang M, Hu Y, Han W, Chen J, Lai J, Wang Y. 2023. Potassium nutrition of maize: uptake, transport, utilization, and role in stress tolerance. The Crop Journal 11: 1048–1058. doi: 10.1016/j.cj.2023.02.009

Zörb C, Senbayram M, Peiter E. 2014. Potassium in agriculture – status and perspectives. Journal of Plant Physiology 171: 656–669. doi: 10.1016/j.jplph.2013.08.008

